# Characterisation of laminar and vascular spatiotemporal dynamics of CBV and BOLD signals using VASO and ME-GRE at 7T in humans

**DOI:** 10.1101/2024.01.25.576050

**Authors:** Sebastian Dresbach, Renzo Huber, Omer Faruk Gulban, Alessandra Pizzuti, Robert Trampel, Dimo Ivanov, Nikolaus Weiskopf, Rainer Goebel

## Abstract

Interpretation of cortical laminar functional magnetic resonance imaging (fMRI) activity requires detailed knowledge of the spatiotemporal haemodynamic response across vascular compartments due to the well-known vascular biases (e.g. the draining veins). Further complications arise from the spatiotemporal hemodynamic response that differs depending on the duration of stimulation. This information is crucial for future studies using depth-dependent cerebral blood volume (CBV) measurements, which promise higher specificity for the cortical microvasculature than the blood oxygenation level dependent (BOLD) contrast. To date, direct information about CBV dynamics with respect to stimulus duration, cortical depth and vasculature is missing in humans. Therefore, we characterized the cortical depth-dependent CBV-haemodynamic responses across a wide set of stimulus durations with 0.9 mm isotropic spatial and 0.785 seconds effective temporal resolution in humans using slice-selective slab-inversion vascular space occupancy (SS-SI VASO). Additionally, we investigated signal contributions from macrovascular compartments using fine-scale vascular information from multiecho gradient-echo (ME-GRE) data at 0.35 mm isotropic resolution. In total, this resulted in *>*7.5h of scanning per participant (n=5). We have three major findings: (I) While we could demonstrate that 1 second stimulation is viable using VASO, more than 12 seconds stimulation provides better CBV responses in terms of specificity to microvasculature, but durations beyond 24 seconds of stimulation may be wasteful for certain applications. (II) We observe that CBV responses show dilation patterns across the cortex. (III) While we found increasingly strong BOLD signal responses in vessel-dominated voxels with longer stimulation durations, we found increasingly strong CBV signal responses in vessel-dominated voxels only until 4 second stimulation durations. After 4 seconds, only the signal from non-vessel dominated voxels kept increasing. This might explain why CBV responses are more specific to the underlying neuronal activity for long stimulus durations.

## Introduction

Cortical layers are a key feature of mesoscopic scale brain organization. Recent developments in functional magnetic resonance imaging (fMRI) have enabled the non-invasive mapping of activation in distinct layer-compartments in humans (Dumoulin et al., 2018). One such technique, Vascular Space Occupancy (VASO, Lu et al., 2003), is a promising cerebral blood volume (CBV)-sensitive signal. In recent years, this method (specifically slice-selective slab-inversion [SS-SI] VASO [Huber et al., 2014], henceforth referred to as VASO) has gained popularity for investigations of cortical depth-dependent signal changes at ultra-high fields (*>*7T) due to its high specificity to the microvasculature and hence the underlying neural activity (e.g. Faes et al., 2023; Haenelt et al., 2023; Huber et al., 2017; Koiso et al., 2023; Persichetti et al., 2020; Pizzuti et al., 2023; Yu et al., 2019; for other VASO variants at ultra-high field, see e.g. Chai et al., 2020; Hua et al., 2013).

The high microvascular weighting of VASO was established by using long stimulus durations (Beckett et al., 2020; de Oliveira et al., 2023; Jin and Kim, 2008), which were also beneficial for mitigating the challenging signal-to-noise ratio (SNR) and sampling efficiency (Huber et al., 2019). However, for neuroscience, the possibility of using short stimuli is often crucial (Huettel, 2012). For example, it allows accounting for carryover effects, enables researchers to study fleeting phenomena such as surprise, or correlate additional factors such as task performance with brain activity. Recently, we have shown that it is feasible to use VASO with short (2 seconds) stimuli and event-related paradigms (Dresbach et al., 2023; for an earlier example of stimuli shorter than 20 seconds, see Persichetti et al., 2020). In this context, we found that stimuli with a duration of 2 seconds had a lower microvascular weighting compared to long stimuli. However, the relationship between (pial) macro-versus (intracortical) microvascular weighting at intermediate, or even shorter stimulus durations is currently unknown for the human VASO signal.

While inferring the micro-or macrovascular weightings of the VASO signal from cortical depth measurements might provide initial insights, detailed information on the vascular architecture would enhance our understanding of signal origins even further. In animal models, this work has been pioneered in the rat somatosensory cortex by various groups. For example, Yu et al. (2016) established intracortical vessel maps by using multi-echo gradient-echo (MEGRE) images and showed that blood oxygenation level dependent (BOLD) and CBV responses originate from venules and arterioles, respectively. On the other hand, Tian et al. (2010) used two-photon microscopy to measure the dilation of individual arterioles and capillaries and showed a backpropagation of arterial vessel dilation. They further compared the microscopy findings to BOLD data acquired with the same stimulation protocol and found good agreement. Although these investigations clearly demonstrate the benefit of combining vascular information with high resolution functional imaging, comparable investigations in humans have been hindered so far by the difficulties of acquiring adequate vascular information in combination with multimodal functional data. Recently, Gulban et al. (2022) showed that acquiring high-quality ME-GRE images is feasible at the mesoscale in living humans. Specifically, the authors provided a scanning protocol that gives high-quality T2* maps acquired at 0.35 mm isotropic resolution and demonstrated that human vascular information can be resolved non-invasively down to the level of intracortical diving vessels (for an alternative approach with time-of-flight magnetic resonance angiography, see Bollmann et al., 2022). Combining this detailed vessel information with high-resolution BOLD and CBV fMRI allows us to investigate the neurovascular coupling in humans.

Therefore, the purpose of this study is twofold. First, we characterized the canonical haemodynamic response function of the layer-dependent VASO signal across stimulus durations common in neuroscience and at high spatiotemporal resolution in humans. Second, we showed a first step in the detailed investigation of submillimeter VASO and BOLD fMRI signals across stimulus durations in vascular compartments by using cutting edge high-resolution anatomical information from ME-GRE data acquired at 0.35 mm isotropic.

## Materials and Methods

### Participants

5 participants without any known neurological condition (3 female, age range: 22-35 years, mean: 26.8 years) took part in the study after providing written informed consent. The study was approved by the Ethics Committee of the Medical Faculty of Leipzig University and the principles expressed in the Declaration of Helsinki were followed.

### Data Acquisition

Scanning was performed in five 90-min sessions per participant (25 sessions in total) using a 7T MRI scanner (Magnetom Terra, Siemens Healthineers, Erlangen, Germany), equipped with a 8Tx, 32Rx RF head coil (NOVA Medical, Wilmington, DE, USA) at the Max Planck Institute for Human Cognitive and Brain Sciences in Leipzig, Germany.

Functional scans were performed in 4 sessions with a 3D echo-planar imaging (EPI) sequence, with SS-SI VASO preparation (Huber et al., 2019; Stirnberg and Stöcker, 2021; VASO version 26dc5a59) and a nominal resolution of 0.9 mm isotropic (16 slices time of inversion [TI]1/ TI2/shot repetition time [shotTR]/echo time [TE] = 1047/2462/42/15 ms, partial Fourier factor = 6/8, variable flip angle scheme with reference flip angle = 33°, 3-fold generalized autocalibrating partially parallel acquisitions (GRAPPA) acceleration [Talagala et al., 2016] in z-direction with a “controlled aliasing in parallel imaging results in higher acceleration” (CAIPIRINHA) shift of 1, bandwidth = 1126 Hz/Px, FoV = 133 mm, matrix size = 148 × 148; (**Figure 1A&D**). B1 shimming was performed in Trueform mode (CP-mode) and the shimming volume included the circle of Willis, to ensure proper nulling of incoming blood. Slice position and orientation were chosen individually and aligned with the left calcarine sulcus (**Figure 1D**, for the functional coverage in all participants, see **Supplementary Figure S1**). To ensure similar slice positions between sessions, we made use of the head scout provided by the vendor.

**Figure 1:**
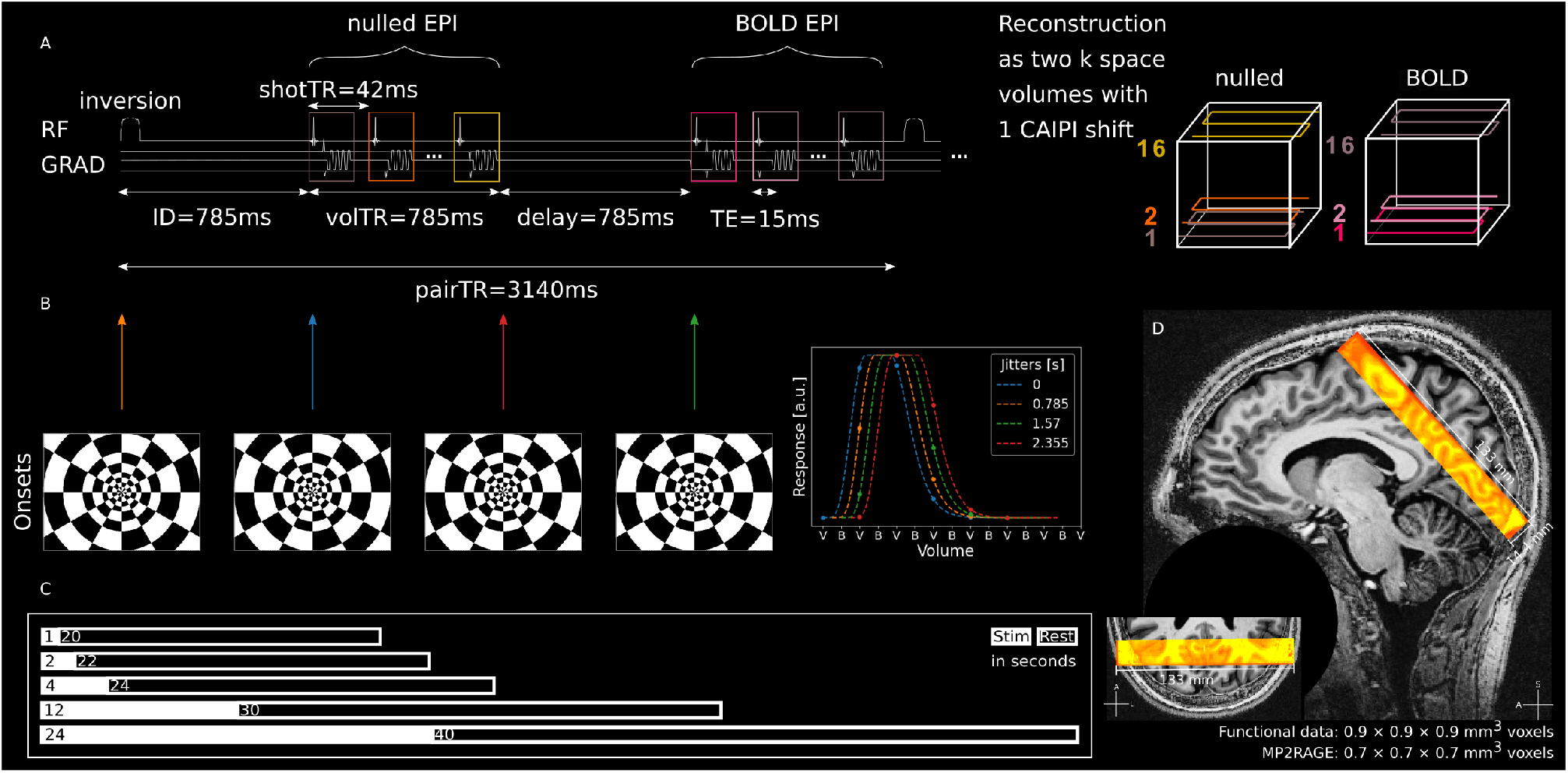
Functional data acquisition and stimulation. **A** Left: Sequence diagram of symmetric VASO acquisition. Right: Illustration if k-space trajectories for nulled and BOLD volumes. **B** Left: Stimulation onsets with respect to acquisition scheme and resulting in subsecond sampling of the stimulus. Right: Illustration of acquired data points of an arbitrary stimulus response per jitter. **C** Stimulus- and corresponding rest durations. Note that in a few sessions (5/20) of individual participants, shorter ITIs were used. For a full description of the scanning sessions, see **Supplementary Table S1**. **D** Coverage of functional slab overlaid on UNI of MP2RAGE (sub-05).

Importantly, we implemented a ‘symmetric’ VASO acquisition, in which an additional delay of 785 ms was inserted between the blood-nulled (further referred as nulled) and BOLD EPI readouts (**Figure 1A**). This delay was as long as the inversion delay and the volume TR, thus creating 4 equally spaced blocks (inversion delay, nulled EPI, additional delay, BOLD EPI) of 785 ms, each, within one acquisition cycle. This had multiple benefits. Firstly, having 4 equally spaced blocks within the acquisition period allowed us to systematically jitter the onset of our stimuli, thereby sampling the haemodynamic response to our stimuli with an effective sampling rate of 785 ms, uncoupling pair TR and effective temporal resolution. Secondly, including the delay before the BOLD EPI acquisition increased the temporal SNR (tSNR) of the BOLD data in previous experiments with a similar setup (Dresbach et al., 2023). Thirdly, the longer T1 relaxation period between inversion pulses led to a better structural T1 contrast in the mean VASO EPI image despite a short volume TR (Dresbach et al., 2023). This was especially important in our case, because it would facilitate the registration of images across sessions.

In a fifth session, we acquired 4 3D ME-GRE images at 0.35 mm isotropic resolution (104 slices; TEs = [3.8, 7.6, 11.4, 15.2, 19, 22.8 ms]; TR = 30 ms, no partial Fourier, flip angle = 12°, GRAPPA factor for in-plane phase encoding = 2; bandwidth = 310 Hz/Px, FoV = 173 mm, matrix size = 496 × 496; volume acquisition time = 13 min, **Figure 6A**). Acquiring 4 images at each echo had 2 purposes. First, we registered and averaged the images to increase SNR. Second, we rotated the phase-encoding orientation by 90°between acquisitions for blood motion artifact mitigation (Gulban et al., 2022; Larson et al., 1990). The slabs of these high-resolution images were centered on the functional coverage (see **Figure 1A** and **Figure 6A**).

Finally, whole-brain images (0.7 mm isotropic, 240 slices, TI1/TI2/TR/TE = 900/2750/ 5000/2.47 ms, flip angle 1/flip angle 2 = 5°/3°, bandwidth = 250 Hz/Px, GRAPPA acceleration factor = 3, FoV = 224 × 224 mm) were acquired in one or two of the sessions using MP2RAGE (Marques et al., 2010). For a full overview of the scanning sessions of all participants, see **Supplementary Table S1**. Scanning protocols are available on github under *<*https://github.com/sdres/neurovascularCouplingVASO/blob/main/protocols*>*.

### Stimulation

In a total of 21-25 runs spread over 4 sessions per participant, flickering checkerboards (contrast reversals at 8 Hz; **Figure 1B**) were presented for 1, 2, 4, 12 and 24 seconds. Stimulus onset was systematically jittered 4 fold with respect to the TR, resulting in an effective stimulus sampling of 0.785 seconds (**Figure 1B**), assuming stationarity of neurovascular responses. Each stimulus duration was presented once per jitter and run, resulting in at least 21 repetitions per participant. Based on a previous study (Dresbach et al., 2023), 20 repetitions per stimulus constitute a good trade-off between efficiency and data stability. Inter-trial intervals (ITIs) were either 10, 12, 14, 20, and 24 seconds (short ITIs) or 20, 22, 24, 30, and 40 seconds (long ITIs, **Figure 1C**). For an overview of which sessions contained runs with short or long ITIs see **Supplementary Table S1**. For sessions with short ITIs, we generated a stimulation design per session and used it for all runs of the session. For the long ITI, we created one stimulation pattern per participant and used it for all runs with long ITIs. We argue that in the latter case, rest durations are sufficiently long to avoid carryover effects. On the other hand, using the same stimulation across more runs allows the averaging of nulled and not-nulled runs across sessions. This might help to reduce noise before dividing nulled by not-nulled time courses during the BOLD-correction. Initially, we had included short ITI sessions for higher efficiency but then opted for long ITIs due to higher signal integrity.

To keep participants’ attention, we implemented a cued button press task. To this end, a gray fixation dot (0.075 degree radius) surrounded by a yellow ring (0.15 degree radius) was presented in the center of the screen. In semi-random intervals, the yellow ring turned into a red target for 500 ms. Participants were instructed to press a button on a button box with their right index finger in a 2 second window after the target. Intervals between targets were chosen randomly between 40 and 80 seconds. The stimulation was controlled with scripts using Psychopy3 (v2022.1.3, Peirce, 2007), which can be found at *<*https://github.com/sdres/neurovascularCouplingVASO/blob/main/code/stimulation/stimulationParadigm.py*>*.

## Data processing and analysis

### MP2RAGE processing

Bias field correction was performed using N4BiasFieldCorrection as implemented in ANTs (Tustison et al., 2010; v2.3.5). Brain extraction was performed based on the second inversion image of the MP2RAGE sequence, using bet (v2.1) as implemented in the FMRIB Software Library (FSL; v6.0.5). The resulting brain mask was applied to the T1w UNI image and the image was further cropped to the region of interest (ROI) to reduce data processing demands. An initial tissue segmentation was performed on the MP2RAGE UNI using FSL FAST (Zhang et al., 2001; v6.0.5). The initial segmentation was manually corrected in the ROI using ITK-SNAP (Yushkevich et al., 2006; v3.8.0) after upsampling to 0.175 mm isotropic using cubic interpolation. Layerification was performed at the upsampled resolution using the LN2 LAYERS program implemented in LayNii (Huber et al., 2021a; v2.2.1) and the equivolume option. For the analysis of temporal data, we estimated 3 layers. To plot layer profiles, we estimated 11 layers.

### Definition of ROIs

All ROIs were defined manually using the macro-anatomical landmark of the calcarine sulcus, indicating primary visual cortex (V1). Specifically, we used the high-quality segmentation and the LN2 MULTILATERATE program in LayNii to grow discs with a cortical parametrization (U,V coordinates), centered on the calcarine sulcus, for each participant. To validate that our ROIs were indeed located in V1, we made use of the fact that V1 is easily identifiable on inflated cortical surfaces and compared the definition to an atlas (Glasser et al., 2016). To define V1 for each participant individually, we followed a standard procedure in BrainVoyager (Goebel, 2012; Version 22.4.4.5188 for macOS) to obtain cortical surfaces. The pipeline is described in detail in the Supplementary Materials. Finally, V1 outlines were drawn corresponding to the definition in the Glasser atlas (Glasser et al., 2016) by author SD and quality controlled by author OFG. Manually drawn outlines of individual participants’ V1 are shown in **Supplementary Figure S2**. To check whether the ROIs fell within the definition of V1, we transformed the resulting surface representation of V1 to voxel space and visually checked the overlap of the V1 definition with our ROIs.

### Functional data processing

The VASO sequence provides two time series. The first consists of ‘nulled’ images, in which the blood signal is as much minimized as possible - ideally 0, giving them a CBV-weighting. The second consists of T2*-weighted BOLD images. If not indicated otherwise, the two time series were treated individually. Data from individual sessions were motion-corrected in a multi-step procedure using ANTsPy (Avants et al., 2011; v0.3.3). First, all volumes of a run were aligned to the first volume of the run using 6 degrees of freedom and rigid body alignment. To guide the registration, we used a brain mask generated automatically using 3dAutomask in AFNI (Cox, 1996; v22.1.13). We then computed run-wise T1w images from the motioncorrected functional data by dividing 1 by the coefficient of variation of the concatenated nulled and BOLD time series on a voxel-by-voxel level. The resulting T1w image of each run was registered to that of the first run of the first session. Finally, the motion correction was performed again by concatenating the transformation matrices of each volume within a run and the transformation matrix generated by the between-run registration. This process was done to minimize consecutive interpolation steps which might introduce unnecessary blurring of the data. Then, the functional images were cropped to the caudal half to reduce computing and storage demands and we averaged all runs within each session. After averaging, we temporally upsampled the data by a factor of 4 to match the stimulus onset grid, duplicated the first volume of the BOLD time series to match the nulled and BOLD data, and divided the nulled by the BOLD data to correct for BOLD-contamination. This resulted in the BOLD-corrected VASO data. Furthermore, we calculated tSNR maps using the LN SKEW program in LayNii and, finally, we now computed a T1w image as before, but now based on the averaged session data to increase SNR.

To estimate spatial stimulus responses, we ran a general linear model (GLM) analysis for the average run of each session using FSL (Woolrich et al., 2001). Here, we used an individual predictor for each stimulation time, convolved with a standard gamma haemodynamic response function without temporal derivative (mean lag: 6 seconds, std. dev: 3 seconds), applied a high-pass filter (cutoff = 0.01 Hz) and no additional smoothing. A second level analysis across sessions was run for each participant. No averaging of statistical maps was performed across participants.

To investigate spatio-temporal responses, we computed voxel-wise event-related averages for each participant. To do so, we averaged responses from the onset of each individual stimulus duration until the end of the respective rest period in each voxel across all sessions. We then computed the baseline as the average activity in the first and last 30 seconds averaged across all sessions. Finally, we calculated % signal changes of the average event-related averages with respect to the average baseline. Note that for VASO, an increase in CBV is shown by negative signal changes. For easier visual comparison with the BOLD data, we inverted the signal changes in all plots.

Due to this averaging procedure, we observed an offset between the time points of the event-related average that were extracted from all runs (until the last volume of the sessions with short ITIs) and the time points that were only extracted from the runs with long ITIs (**Supplementary Figure S3**). To account for this, we subtracted the difference in signal change between the last time point of the short ITI sessions and the first time point of the long ITI sessions from the time points only present in the long ITI sessions. This results in two time points with the same value for each participant and stimulus duration, which is not visible in the group data. Finally, we set the value of the first time point of a given event-related average to 0 and normalized the remaining time points to this. This procedure was done for each participant, layer and stimulus duration independently before averaging across participants (For a graphical representation, see **Supplementary Figure S3**).

### ME-GRE data processing

ME-GRE data were processed as described in Gulban et al. (2022). Key points of the analysis are the blood motion artifact mitigation and the correction of head motion between acquisitions. The blood motion artifact is a displacement of signal from arterial vessels due to the read phase encoding direction (Larson et al., 1990). This issue was mitigated by acquiring data with four different phase encoding directions and compositing new images with less influence of the artifact. To correct for head motion between acquisitions, we registered the averaged echoes of individual runs to the first run and applied the resulting transformation matrix to the individual echoes. In this process, we also upsampled the images to 0.175 mm isotropic using cubic interpolation and finally averaged images from four acquisitions to boost SNR. From the resulting images, we calculated T2* and S0 values. Based on the T2* data, we performed a manual vessel segmentation in ITK-SNAP restricted to our ROI. Specifically, we selected low-intensity voxels arranged in tube-like structures (**Figure 6B**).

### Registration

The statistical maps from the functional data analysis and the vessel segmentation based on the ME-GRE data were registered to the upsampled anatomical MP2RAGE data. To register the functional data, the upsampled UNI images of the MP2RAGE data were first registered to the T1w image in EPI space derived from all sessions. The registration was performed using the cross-correlation metric and the non-linear SyN algorithm of antsRegistration as implemented in the ANTS toolbox (Avants et al., 2011; v2.3.5) to account for the geometric distortions in the functional data. Furthermore, the brain mask generated during the motion correction was used to guide the registration. The resulting linear and non-linear transformation matrices were then used to inversely register the statistical maps and event-related averages to the anatomical MP2RAGE data at upsampled resolution (0.175 mm isotropic).

To register the ME-GRE data, the upsampled MP2RAGE inv-2 images were registered to the upsampled S0 images resulting from the ME-GRE data. Here, we first manually matched the images using ITK-SNAP. This initial alignment was then improved by registering the images using ITK-SNAP with the mutual information metric and a mask covering the posterior brain border generated during the ME-GRE processing pipeline. Finally, we used the inverse registration matrix to register the vessel segmentation to the anatomical data.

## Results

### Behavioral performance, functional data quality, and head motion

To keep participants’ attention during the functional sessions, they performed a cued button press task. Analysis of the attention task revealed that participants consistently performed well (**Supplementary Figure S4A**). Only in 3 runs, the ratio of detected/total targets was *<*0.75. Given that targets could be presented alongside the task stimulus and therefore hard to detect, we did not exclude any data based on isolated undetected targets. We further examined how many consecutive targets were missed within runs, as missing multiple targets in a row could be indicative of participants falling asleep. We found that in 2 runs 2 targets, and in two other runs 3 consecutive targets were not detected. Given this low number and an average of 11 targets per run, we did not exclude data based on attention task performance.

Participant head motion is a common problem in fMRI, especially for high resolution data. To assess head motion, we computed framewise displacements (FDs) from the rigid motion parameters given by the motion correction (Power et al., 2012). The distribution of volumes which were displaced with respect to the previous volume by more than the voxel size (0.9 mm) is shown for nulled and BOLD timecourses separately in **Supplementary Figure S4B**. Only a small number of volumes (92; *<*0.08 %) showed FDs greater than our voxel size (0.9 mm). 40 of those (in 16 individual runs) were in BOLD and 52 (in 12 individual runs) were in nulled time series and FD never exceeded 1.88 mm (2*×* voxel size). We therefore did not exclude any data based on motion.

**Figure 2A** shows the high data quality in a representative participant (sub-06) after averaging data from an individual session. Both mean nulled and BOLD echo planar images look crisp with minimal artifacts. Note the outstanding anatomical contrast in the mean nulled images. This helped in the correction of motion within runs and especially the registration between runs and sessions.

**Figure 2:**
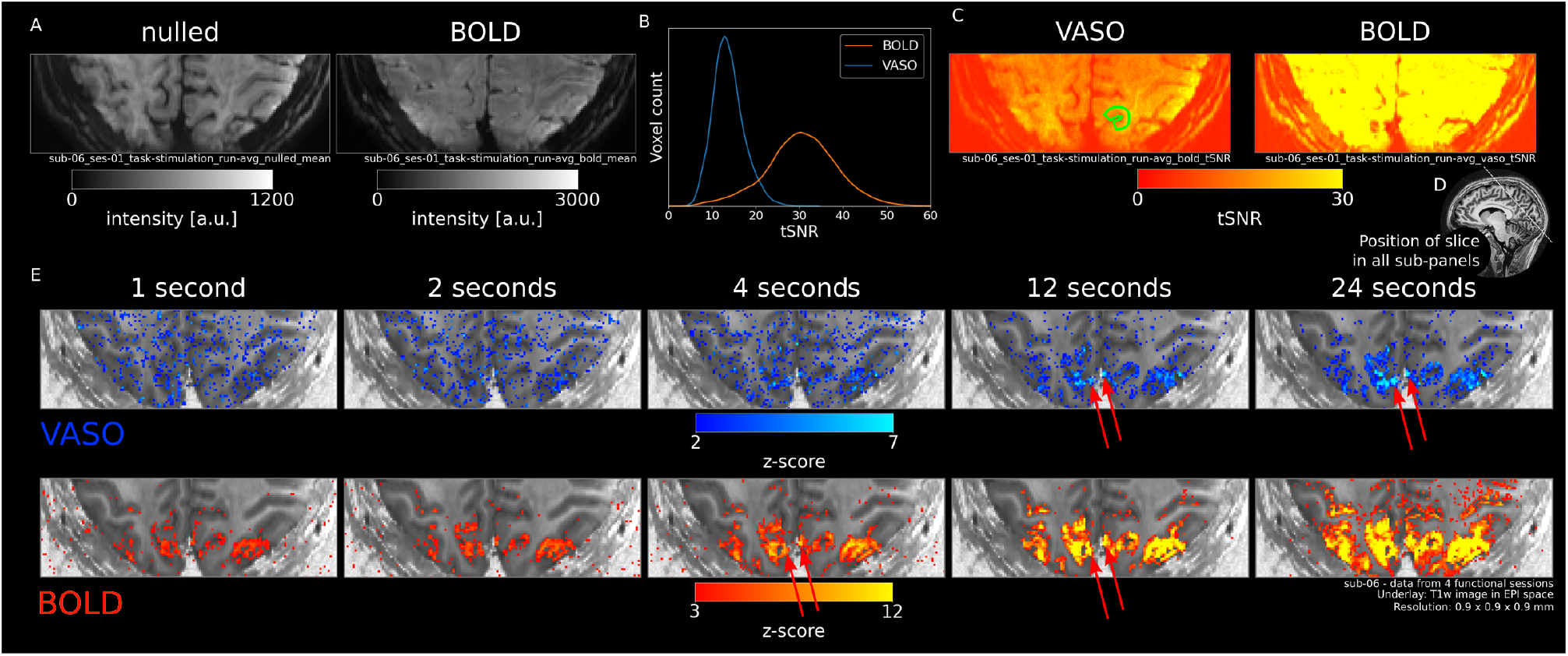
Quality assessment and activation. **A** Mean nulled and BOLD images of all runs in the first session (n=6) of a representative participant (sub-06). **B** BOLD and VASO tSNR across all participants extracted from ROI (see **Figure 3A** and outline on a single slice in **Figure 2C**). **C** VASO (left) and BOLD (right) tSNR maps of one participant (sub-06) averaged across all runs of the first session (n=6). Green outline shows the ROI on the slice. Note that ROIs extend across multiple slices (see Figure 3A). **D** Position of the slice shown in all sub-panels illustrated on a whole-brain MP2RAGE UNI image. **E** VASO (upper) and BOLD (lower) z-maps of one participant (sub-06) resulting from GLM (contrast: stimulus duration rest) for all stimulus durations separately, averaged across all sessions. Red arrows indicate likely position of large pial vessels.

**Figure 2B** shows the tSNR across participants for VASO and BOLD within the ROIs (for an example ROI, see **Figure 3A**). tSNR values were well above 10 for VASO and above 30 for BOLD. For tSNR values of individual participants and sessions within the ROI, see **Supplementary Figure S5**. Individual tSNR values are mostly uniform across sessions with minor differences between participants. **Figure 2C** shows an example of VASO and BOLD tSNR maps of one individual participant (sub-06) after averaging one session.

**Figure 3:**
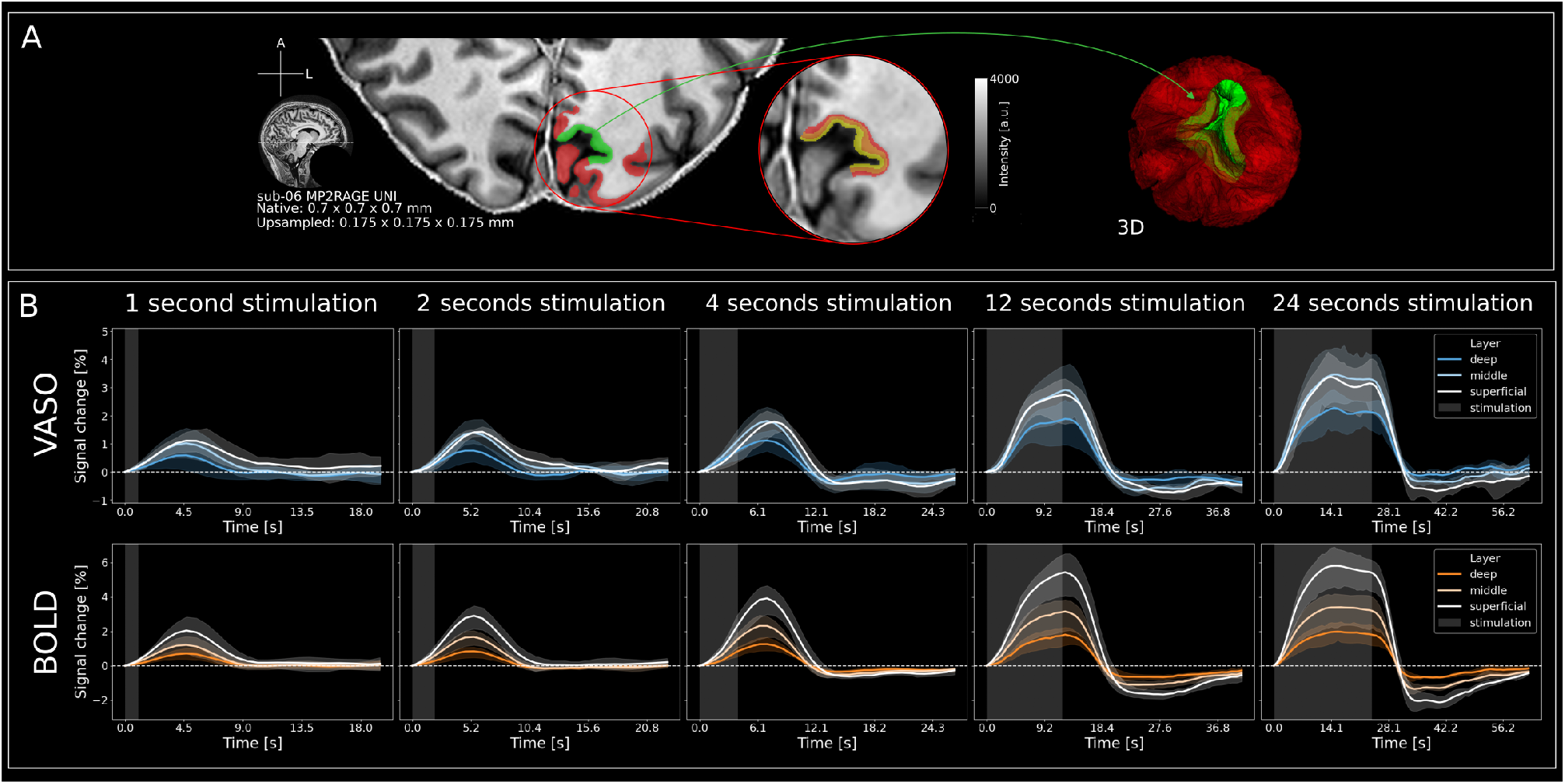
Segmentation and event-related averages across cortical depth. **A** Left: Cortical gray matter segmentation (red & green) overlaid on a single slice of the MP2RAGE UNI image of one participant (sub-06). The ROI in the calcarine sulcus (V1) is indicated by green color. Middle: Zoomed in section of the segmentation in ROI with layering. Right: 3D-representation of the segmentation. Red translucent shape is general gray matter while green refers to the ROI. Note that here, only 3 layers are shown. For the layer profiles in **Figure 4**, we estimated 11 layers. **B** Group level (n=4) VASO (upper) and BOLD (lower) signal changes [%] in response to all stimulus durations separately for superficial, middle and deep layer compartments. Note the subtle differences of the response time, magnitude, and HRF shape across layers, stimulus durations and contrasts. Stimulation period is indicated by gray shaded areas and error bands indicate 95% confidence interval across participants.

Taken together, we concluded that the functional data was of sufficient quality to proceed with further analyses.

Obtaining high quality statistical maps is often difficult in layer-fMRI. Therefore, investigating voxel-wise results from the GLM analysis constitutes a rigorous assessment of data quality. For VASO, longer stimuli (12 seconds) evoked clear responses in visual cortices (**Figure 2E**, top) showing the high data quality. Notably, the activation is centered in gray matter, as expected based on the high specificity of CBV measurements. However, we also observed sparse occurrences of highly active regions for VASO in cerebrospinal fluid (CSF), likely reflecting a dilation of pial vessels (red arrows in **Figure 2E**). While in response to 4 seconds stimulation, some activation can be found in gray matter for VASO, only weak to no visible patterns can be observed for stimuli with a duration of 1 or 2 seconds. For BOLD, all stimulus durations lead to clear responses whose amplitudes increased with stimulus duration (**Figure 2E**, bottom). As expected for BOLD, strongest responses can be observed in CSF due to the draining vein effect. Taken together, the results from the statistical maps are in line with our expectations due to the limited number of trials and low tSNR of VASO compared to BOLD (**Figure 2B/C**).

### Temporal stimulus evoked responses across layers in V1

To quantify the signal changes across cortical depth and time, we performed a high quality semi-manual segmentation of cortical gray matter (**Figure 3A**). We then extracted event-related averages from 3 cortical layers in V1. Specifically, we computed a geodesic cylindrical ROI centered in the calcarine sulcus using LN2 MULTILATERATE as implemented in LayNii (green and layered shapes in **Figure 3A**). The responses of individual participants are shown in **Supplementary Figure S6** for VASO and **Supplementary Figure S7** for BOLD. Most participants showed clear stimulus evoked responses for all stimulus durations and layer compartments for both VASO and BOLD. However, we did not observe reliable VASO activation in response to stimulation of 1 and 2 seconds for one participant (sub-08, see **Supplementary Figure S6**). This participant was therefore excluded from further analyses. Group-level (n=4), time-resolved VASO and BOLD responses for 3 cortical depths and all stimulus durations are shown in **Figure 3B**.

Most notably, VASO has slightly stronger responses in superficial compared to middle and deep layers for short stimuli (*≤*4 seconds). On the other hand, longer stimulus durations evoked slightly stronger VASO responses in middle layers, which became more pronounced with increasing stimulus duration. For a more conventional representation of depth-dependent activity, we extracted laminar profiles from VASO and BOLD z-maps resulting from a GLM analysis with a predictor for each stimulus duration contrasted against the rest periods (**Figure 4**). For VASO, the results show clear peaks in middle layers for all stimulus durations (**Figure 4A** left), whereas BOLD peaks are located in superficial layers (**Figure 4B** left). To show differences in profile shape irrespective of activation amplitude, we min-max normalized the responses across cortical depth (**Figure 4A&B** right). Also here, it can be seen that short stimulus durations show a slightly stronger response in superficial layers than long stimuli for VASO.

**Figure 4:**
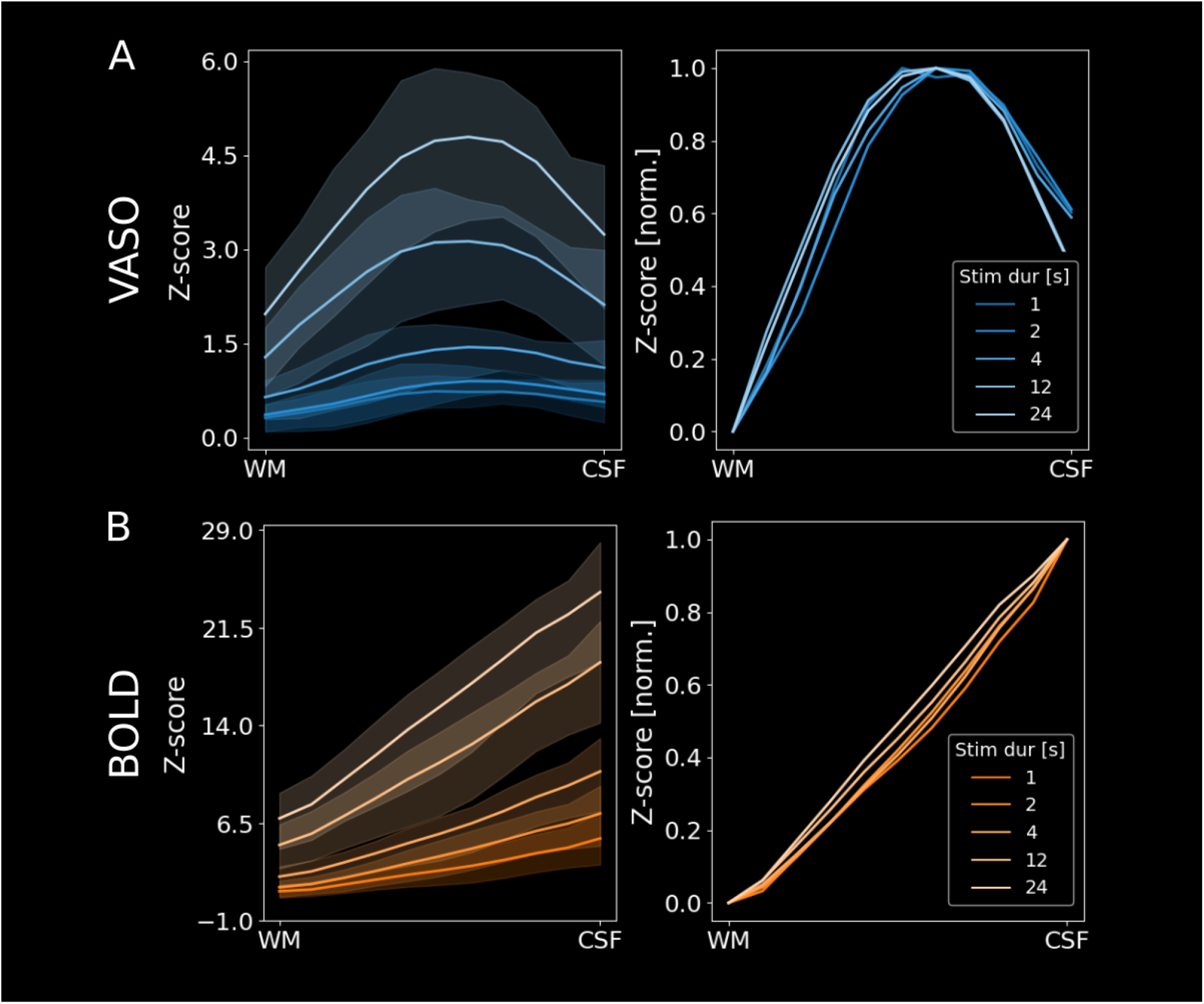
Depth-dependent profiles show clear peaks in middle cortical layers and modulation of microvascular weighting for VASO. **A** Left: Group level (n=4) VASO layer profiles showing z-scores across cortical depth for all stimulus durations separately. It can be seen that the VASO response decreases less strongly in superficial layer groups for shorter stimulus durations. Error bands indicate 95% confidence intervals across participants. Right: Same data as on the left, but min-max normalized to show differences in profile shape irrespective of activation amplitude. It can be seen that the VASO response has a relatively stronger weighting towards superficial layer groups for short stimuli (more concave). Error bands were omitted for better visibility because of the large overlap. **B** Same as **A** but for BOLD data.

Note, that the clearer middle layer peaks for all stimulus durations in VASO based on the layer profiles (**Figure 4A**) compared to the event-related averages (**Figure 3B**) stems from the fact that the former are based on z-scores whereas the latter are based on raw % signal change. As superficial layers usually exhibit higher noise levels compared to middle or deep layers, responses close to the cortical surface are attenuated when the signal is plotted in terms of z-scores. For completeness, z-scored event-related averages are shown on the group level in **Supplementary Figure S8** for VASO and BOLD (see **Supplementary Figure S9** and **Supplementary Figure S10** for individual participants’ z-scored VASO and BOLD event-related averages, respectively). Here, middle layer activity for VASO is clearly strongest for all stimulus durations, resembling the layer profiles.

Remaining apparent differences between event-related average and layer profile representations stem from the higher partial voluming when extracting signals from 3 (as done for the event-related averages) rather than 11 layers (as done for the layer profiles). For BOLD, response-strength increases from deep to superficial layers, irrespective of stimulus duration, or form of representation, as expected based on the draining vein effect.

We also find delayed VASO peaks in superficial layers compared to middle and deep layers for short stimuli (**Figure 3B**). To quantify this delay, we defined the time to peak (TTP) as the time between stimulus onset and the time point of highest signal change for each stimulus duration and layer compartment separately. The results for VASO and BOLD are shown in **Figure 5**. For VASO, there are 3 noteworthy observations. Firstly, as described based on visual assessment above, the TTP was the highest in superficial layers for short stimuli. Secondly, the highest TTP is located in the middle layers for long stimuli. Thirdly, we find that for stimuli up to 12 seconds, the response peaks after stimulus cessation for all layers, while the response to 24 second stimulation drops off before the stimulation ends. For BOLD, the response to stimuli with a duration of 24 seconds also peaks before stimulus cessation and shows the longest TTP in middle layers.

**Figure 5:**
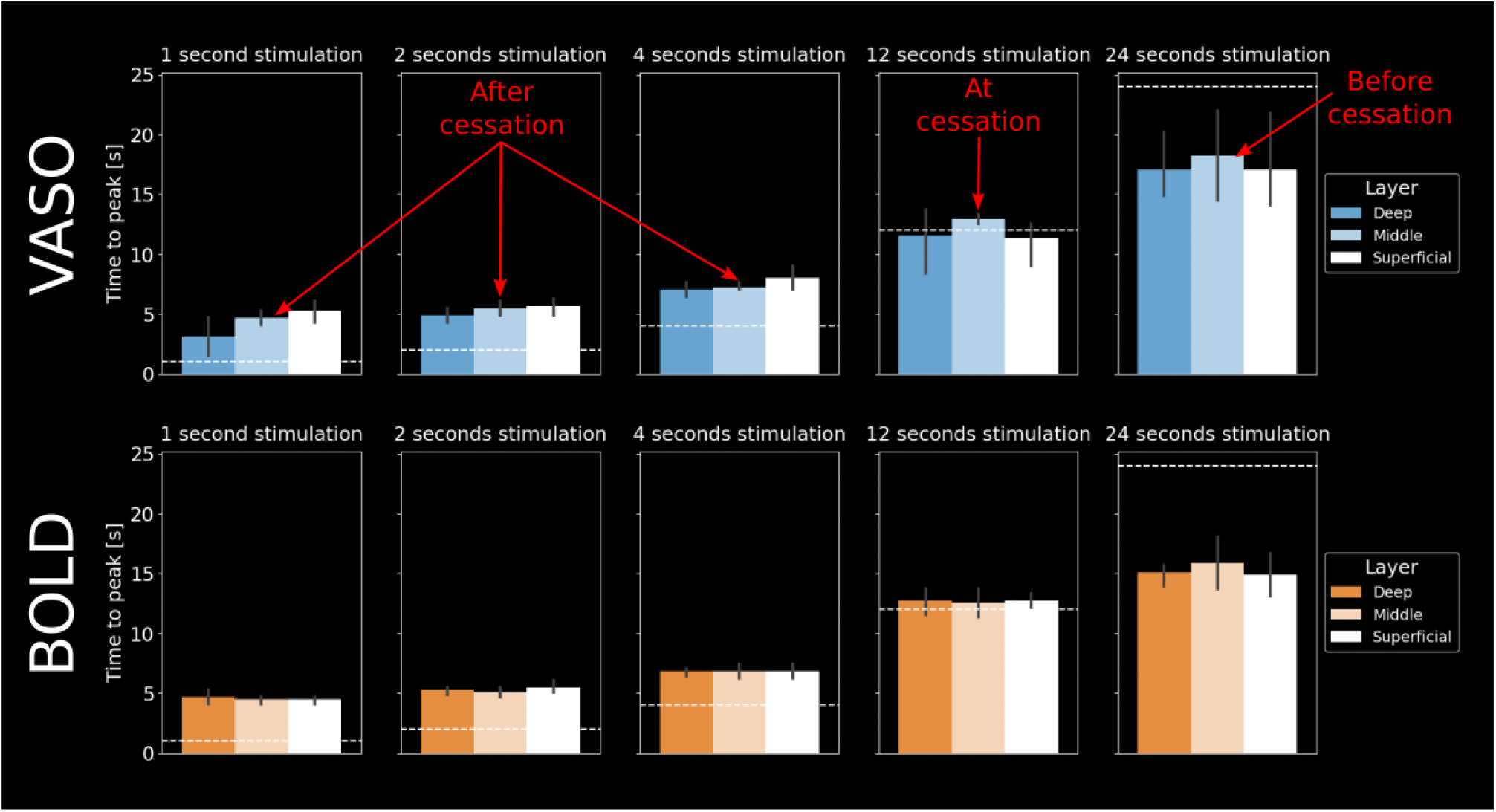
Time-to-peak analysis. VASO (upper) and BOLD (lower) time to peak (TTP) for all stimulus durations (indicated by white dashed line) and layers independently. For short stimuli, the TTP is after stimulus cessation. For stimuli with a duration of 12 seconds, the TTP is around stimulus cessation. For stimuli with a duration of 24 seconds, the TTP is after stimulus cessation. Furthermore, it can be seen that the layer-specific distributions of TTPs change across stimulus durations. For short stimuli, the superficial layers peak later, whereas for longer stimuli the middle layers tend to peak later. This could be explained by different relative weighings of various vascular compartments (see **Figure 6**). For BOLD, we could not see any trends of layer-dependent TTPs across any stimulus durations. TTP was defined as the highest signal change between the initial deflection above 0 and falling off below 0. Error bars indicate 95% confidence intervals across participants.

**Figure 6:**
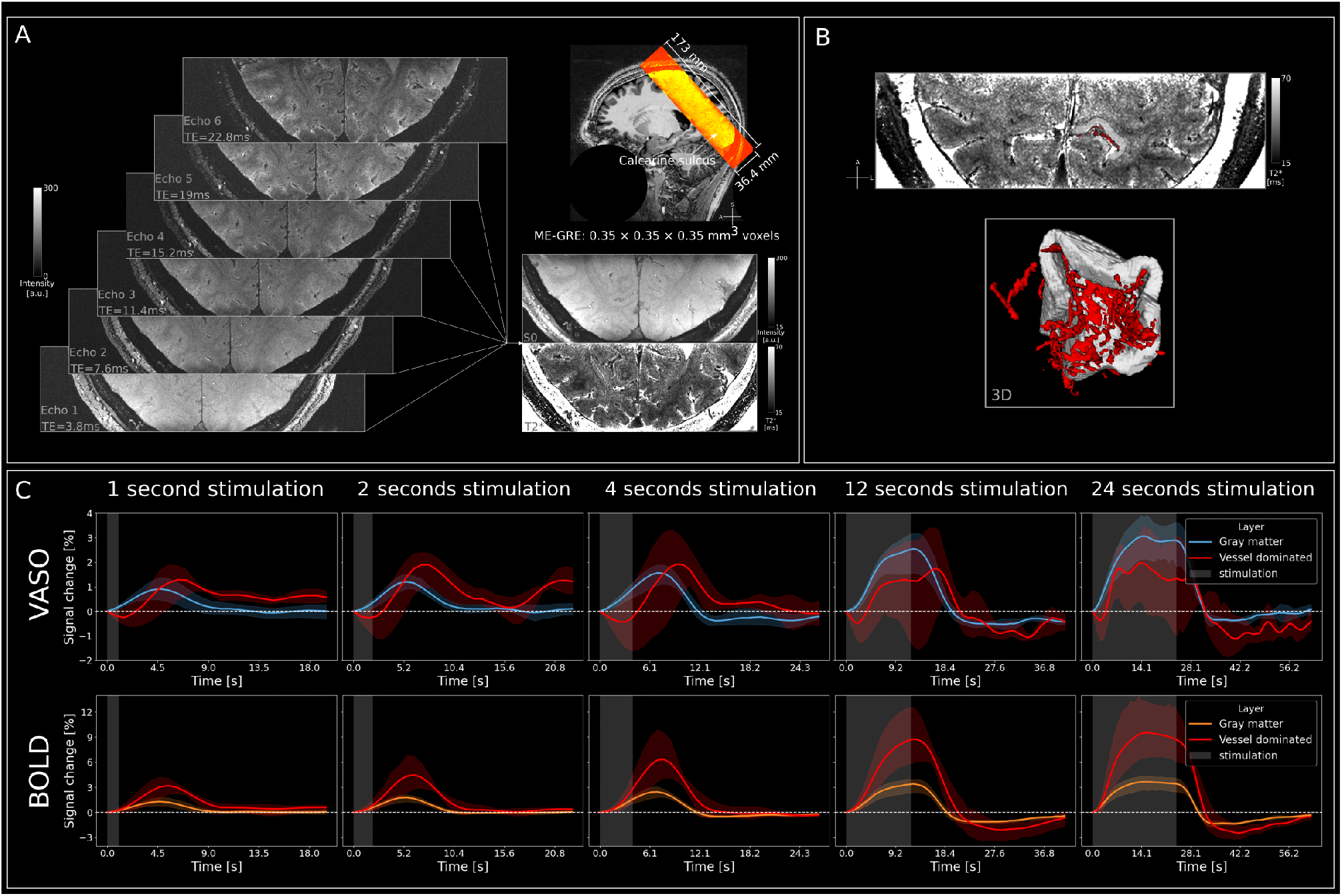
Signal in vessel dominated voxels shows additional signal dynamics. **A** Acquisition of ME-GRE data at 0.35 mm isotropic resolution. Acquired slab overlaid on the MP2RAGE UNI image (sub-05) **B** Quantitative T2* map and segmentation of vessels and ROI in a representative participant (sub-06, top) and a 3D representation of the same data (bottom) **C** Group Level (n=4) VASO (top) and BOLD (bottom) signal changes in percent for gray matter and vessel dominated voxels separately for each stimulus duration. It can be seen that for short stimuli VASO shows slightly larger responses in large vessels compared to gray matter. For longer stimuli, parenchyma responses are stronger than vessel responses. BOLD shows strong vessel responses, markedly exceeding gray matter across all stimuli. Stimulation period is indicated by gray shaded areas and error bands indicate 95% confidence interval across participants.

### VASO and BOLD signal in vessel dominated voxels

In order to investigate the origin of VASO and BOLD responses, we computed signal changes in vascular structures compared to gray matter. For this, we manually delineated vessels based on the ME-GRE data **Figure 6B**, which can be for example identified by voxels with very short apparent T2* values (due to strong dephasing inside the blood) and an arrangement in a tube-like fashion. **Figure 6C** shows the group level (n=4) responses for gray matter and vessel dominated voxels. Data from individual participants are shown in **Supplementary Figures S11** and **S12** for VASO and BOLD, respectively. There are several noteworthy observations. Firstly, the vessel-dominated VASO signal change shows delayed peak times compared to signals originating predominantly from gray matter for all stimulus durations up to and including 12 seconds (**Figure 6C**). This could explain the delayed peak in superficial layers of gray matter as observed in **Figure 3B**. Secondly, for short stimuli, the superficial vessel response is stronger than that of gray matter. On the other hand, gray matter VASO signals dominate for stimuli of 12 and 24 seconds. This is driven by a slow increase of VASO gray matter activity from short to long stimulus durations, while the response in vessel dominated voxels is saturated after 2 seconds of stimulation and does not rise even for 24 second stimuli. For BOLD, vessel dominated voxels show a markedly larger response than gray matter, irrespective of the stimulus duration, with responses that keep increasing for longer stimuli. **Figure 7** illustrates the ratio between macrovascularly dominated and gray matter peak-signal changes for both VASO and BOLD, separately for each stimulus duration. For VASO, the data show again that only for long, and not short stimuli, the gray matter response is larger than the response in macroscopic vessels. For BOLD, the data show that the relationship between BOLD activity in vessels and gray matter stays very similar, irrespective of stimulus duration.

**Figure 7:**
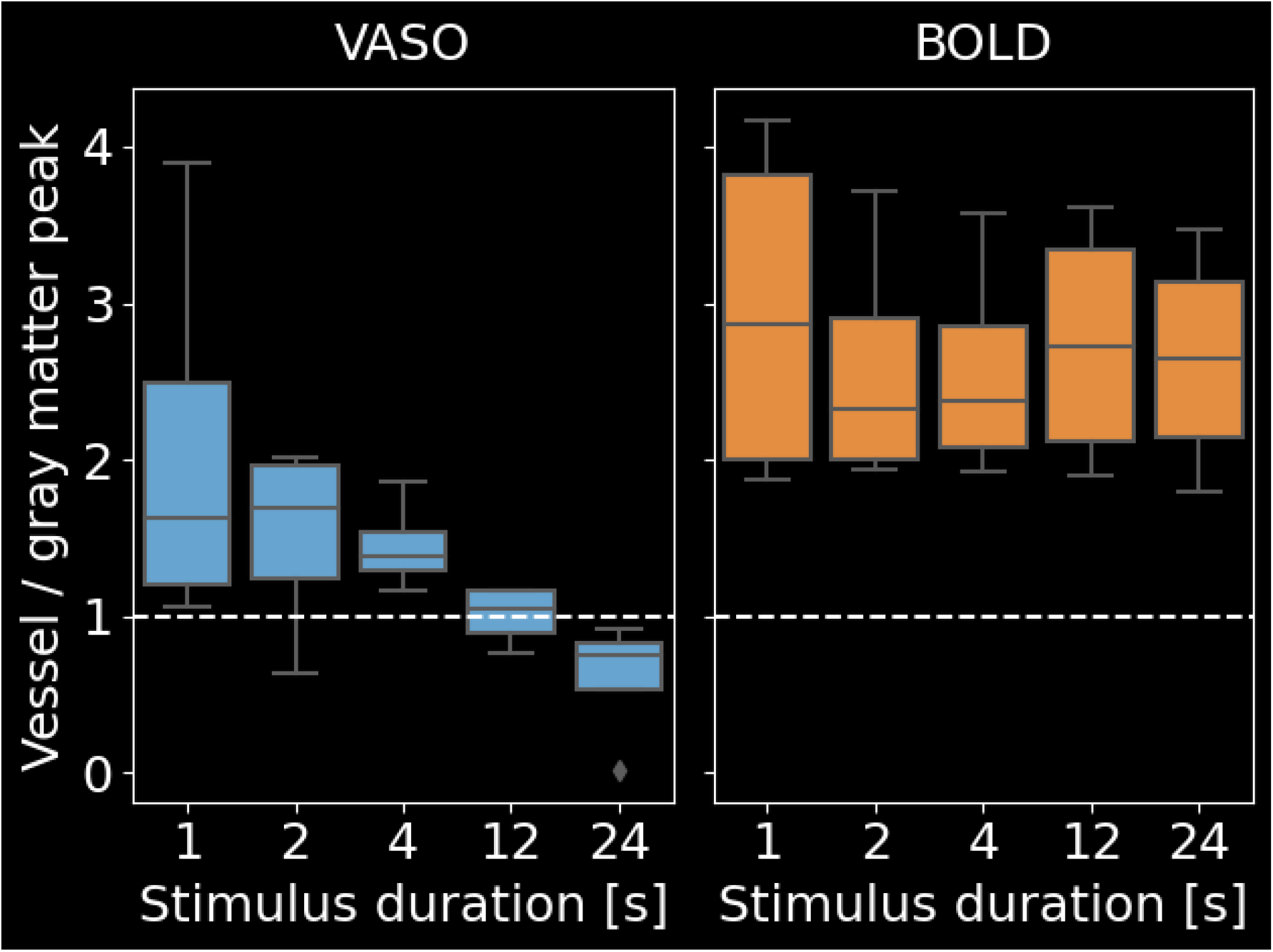
Signal in vessel dominated voxels shows additional signal dynamics. VASO (left) and BOLD (right) peak signal change of vessel dominated voxels divided by the peak signal change in gray matter for each stimulus duration independently. For VASO, the ratio depends on the stimulus duration, whereas for BOLD, this influence is negligible. Overall, the vessel bias is smaller for VASO than for BOLD across stimulus durations. It can be seen that for VASO, long stimuli result in responses that are more confined to gray matter.

## Discussion

### Summary of results

In the present study, we characterized laminar CBV responses to visual stimuli with a range of durations that are common in neuroscientific investigations. To do so, we used high spatiotemporal resolution (effective temporal sampling: 785 ms; spatial resolution: 0.9 mm isotropic) VASO fMRI, in combination with cutting-edge anatomical ME-GRE data acquired at 0.35 mm isotropic to glean vascular information. With these data, we showed that the microvascular weighting of the VASO signal depends on stimulus duration and could attribute this effect to the ratio between activity in gray matter and superficial, vessel-dominated voxels.

### Implications for neuroscientific applications

Our results have several implications for neuroscientific applications of layer-specific VASO, especially with respect to the stimulus duration. Firstly, we were able to obtain reliable responses to stimuli with a duration of 1 second in 4 out of 5 participants. After Persichetti et al. (2020) with a stimulus duration of 6 seconds and Dresbach et al. (2023) with a stimulus duration of 2 seconds, to our knowledge, this is the shortest stimulation duration with which VASO responses have been seen at submillimetre resolution in humans. For neuroscientific purposes, shorter stimulus durations are often desired, as they, for example, allow to investigate transitory phenomena like surprise or enable better modulation of stimulus detectability in cases where detection at threshold is desired (Huettel, 2012). Therefore, our results may facilitate future investigations of novel research questions using laminar-specific VASO.

However, the shorter stimulation time seems to come at the cost of spatial specificity to the underlying neural populations. Specifically, we further showed that the laminar weighting of VASO differed between stimulus durations. Shorter stimulation durations (*≤*4 seconds) showed highest % signal changes in superficial layers, whereas during longer stimulation (*≥*12 seconds) signal changes in middle layer activity were highest (**Figure 3B**). Note that in contrast to raw % signal changes, the peak z-scores were in middle layers for all stimulation durations (**Figure 4A**), however with a more shallow decrease towards the cortical surface for short stimuli. This can be explained by the higher noise level in superficial layers which downscales the response at the cortical surface when interpreting the data in terms of z-scores (for z-scored event-related averages, see **Supplementary Figures S8-S10**). These findings are in line with our previous VASO results comparing block-to event-wise stimulation in humans (Dresbach et al., 2023) and animal work using MION contrast agent (e.g. Jin and Kim, 2008). However, in our previous study, we used either short (2 second), event-related or long (30 seconds), block-wise stimulation and could not determine the stimulation duration at which the relationship between micro- and macrovascular weighting changes is optimal. Based on the present study, stimulus durations of 12-24 seconds or longer might be necessary for the VASO signal to develop the maximal weighting towards the microvasculature. However, the gap between the 4 second stimulus and the 12 second stimulus studied here is still substantial. Therefore, future studies could investigate whether other durations within this range are sufficient to develop the desired spatial specificity.

Finally, we found that for stimuli up to and including 12 seconds, the VASO TTP was after stimulus cessation (**Figure 5**). However, for 24 second stimuli, the response reached its highest point before the stimulus ended. Note that the TTP was longest in superficial layers for short stimulation durations, probably due to the largest contribution of larger vessels that react slightly slower than the microvasculature close to the neuronal tissue in gray matter. On the other hand, TTP was longest in middle layers for longer stimuli, hypothetically due to the slow, passive dilation of the capillary bed. However, this effect is rather small and is best interpreted with caution.

Taken together, several factors need to be considered when choosing the stimulus duration in future VASO experiments. Based on our experiments, longer stimuli (*≥*24 seconds) are advisable for higher signal amplitudes, superior detection sensitivity and good microvascular weighting. On the other hand, to capture transient brain activity dynamics, shorter stimuli are better suited. Finally, if one is interested in obtaining the highest possible magnitude of the response while being temporally efficient, stimuli with a duration between 12 and 24 seconds might be advisable.

### Superficial vessel contributions in BOLD and VASO

In line with various studies using BOLD as measured with gradient-echo (Bause et al., 2020; Kay et al., 2019; Kemper et al., 2018; Moerel et al., 2018), spin-echo (Koopmans and Yacoub, 2019), or 3D-GRASE (Moerel et al., 2018), we find that BOLD signal changes are much larger in vessel dominated voxels than in gray matter. Furthermore, our results show that the relationship between BOLD signal changes in vessel dominated voxels and gray matter does not change substantially across stimulus durations **Figure 7**. As a result, the specificity of BOLD responses seems to be somewhat independent of the duration of stimulation. For VASO, we also find signal changes in vessel dominated voxels. VASO responses in pial vessels have been highlighted in the past by us and other groups in numerous conference abstracts (Handwerker et al., 2016; Huber et al., 2015; Venzi et al., 2019), user manuals (Huber, 2018a, 2018b), and in a previous paper (Huber et al., 2021b). However, they have never been systematically investigated. In the following we will discuss three main observations.

Firstly, we find consistently delayed onsets of vessel dominated compared to gray matter VASO signal changes for all stimulus durations. This matches our previous results in humans (Dresbach et al., 2023) and results from Tian et al. (2010) who found earliest dilation of capillaries in middle gray matter, followed by a backpropagation towards the cortical surface and later activation of larger feeding vessels in rats, using two-photon microscopy.

Secondly, we find consistently delayed peaks of vessel dominated compared to gray matter VASO signal changes for short stimuli only. In a previous study (Dresbach et al., 2023), we found similarly delayed VASO response peaks of superficial, compared to middle and deep layers. Potentially, the current finding indicates that this effect could have been (in part) driven by partial voluming with pial vessels. However, we cannot exclude a gradual process in line with the serial backpropagation proposed by Tian et al. (2010).

Finally, we find that vessel dominated activity saturates after 2-4 seconds of stimulation. This large initial VASO response in superficial vessels may result from the actively muscle-controlled dilation of superficial arteries, which reaches its maximum already after short stimulation times (Kennerley et al., 2012). On the other hand, we found that gray matter VASO activity continued to rise with stimulus duration, potentially through the slow and passive dilation of the capillary bed. As a result, the ratio of vessel-dominated over gray matter VASO activation is variable, with highest gray matter weighting for longer stimuli.

In summary, this leads to three phases of the VASO response. An early gray matter response that is mostly free of superficial vessel activity for all stimulus durations. Although the signal is small, this phase is in theory very specific to the underlying neuronal activity as it is not contaminated by pial vessel dilation. This could lead to interesting applications of fast imaging protocols with short stimulation and highly jittered paradigms using VASO. In the second phase, activation is mostly dominated by larger vessels. This is the least specific phase, as the macrovascular component exceeds that of gray matter. In the third phase, specificity is high again with larger gray matter than macrovascular weighting.

### Limitations & Future directions

As discussed in the previous section, we have explored the combined use of high resolution functional VASO data with ME-GRE data at 0.35 mm isotropic to investigate CBV and BOLD responses in superficial vessel dominated voxels - so far without differentiation between arterial and venous contributions, as this was outside the scope of the current investigation. Importantly, previous studies in animals were able to differentiate (intracortical) arteries from veins and showed that CBV responses mainly originate from the arterial vasculature, whereas BOLD responses mainly originate from venous vasculature (Bolan et al., 2006; Chen et al., 2019, 2021; Yu et al., 2012, 2016). Therefore, future studies could investigate the contributions of superficial and intracortical arteries and veins to the CBV and BOLD signal acquired with VASO in humans.

Identifying signal changes in the vasculature is, however, hindered by the functional spatial resolution of 0.9 mm isotropic employed in the present study. Specifically, superficial cortical arteries and veins were found to have diameters up to 280 and 350 *µ*m, respectively (Duvernoy et al., 1981).^1^ Furthermore, intracortical vessels were found to have diameters up to 125 *µ*m and have been shown to be organized in vascular units with central veins (separated by approximately 1-1.5 mm), surrounded by rings of smaller arteries (Bolan et al., 2006; Duvernoy et al., 1981; Harel et al., 2010). Assuming that the arterial rings are roughly centered between two veins, this would yield an artery-vein distance of about 0.5-0.75 mm. As a result, both the superficial and intracortical vessels are significantly smaller and/or closer together than we can resolve with our voxel size of 0.9 mm isotropic. Therefore, partial voluming effects have to be taken into account for functional responses from superficial vessels investigated in the present study, and will limit the distinction between signal changes originating from arteries and veins in future investigations.

Finally, the VASO signal is usually optimized to obtain functional responses in gray matter and the sequence protocol used here is exceptional in various ways; potentially leading to inflated signal changes above the cortical surface. For example, the use of very short TEs (15 ms) and incorporation of data from multiple sessions may lead to mis-corrections of BOLD contaminations in large superficial vessels due to BOLD-overcompensation and potential misregistration (Chaimow et al., 2019; Devi et al., 2022; Genois et al., 2021; Huber, 2014, 2018a). Furthermore, the strong, (flip angle dependent) contrast between CSF and gray matter in our data may lead to apparent signal changes in superficial vessels due to volume redistributions upon activation (Donahue et al., 2006; Jin and Kim, 2010; Piechnik et al., 2009). Finally, the relatively short TR may lead to CBF-dependent outflow of not-inverted water into the post-capillary vasculature, resulting in lower gray matter signals (Wu et al., 2010) and incomplete nulling of draining veins (Huber, 2014 - Thesis chapter 5). Taken together, future studies with shorter experiments to avoid the integration of multiple sessions or different combinations of TEs, flip angles and TRs will help with the interpretation of the presented results and with determining how far they generalize.

## Conclusion

We characterized the spatiotemporal dynamics of CBV and BOLD responses across cortical depth and vessel dominated voxels at 0.9 mm spatial, 0.785 seconds effective temporal resolution and *>*7.5h of (f)MRI scanning per participant. We demonstrated that it is feasible to obtain reliable VASO responses to stimuli with a duration of 1 second and showed differences in vascular weighting of CBV signal changes depending on stimulus durations, which are fundamental to future neuroscientific applications of VASO. Furthermore, we found differences in signal dynamics across time, space and stimulus duration for VASO and BOLD, which might inform laminar models of the haemodynamic response. Finally, we found strong VASO signal changes in superficial vessel-dominated, compared to gray matter voxels for short stimuli and stronger gray matter activity for long stimuli, which gives fundamental insights into mechanisms of the VASO contrast.

## Acknowledgments

We thank Domenica Klank and her team of medical technical assistants at the Max Planck Institute for Cognitive and Neuroscience in Leipzig for their help with participant handling, booking and general assistance while scanning. We thank Rüdiger Stirnberg and Tony Stöcker from DZNE in Bonn for sharing, optimizing, and supporting us with the 3D-EPI VASO sequence used here. We thank Vojťech Smekal, Till Steinbach and Maite van der Miesen for helpful discussions on the manuscript. Special thanks goes to the remaining “Maastricht layer-seminar” members Lonike Faes, Lasse Knudsen and Kenshu Koiso, for countless discussions on topics related to layer-fMRI. Finally, we thank Kamil Uludag for the inspiring discussions on neurovascular coupling.

SD is supported by the ‘Robin Hood’ fund of the Faculty of Psychology and Neuroscience and the department of Cognitive Neuroscience. RG is partly funded by the European Research Council Grant ERC-2010-AdG269853 and Human Brain Project Grant FP7-ICT-2013-FET-F/604102. OFG is funded by Brain Innovation. NW has received funding from the European Research Council under the European Union’s Seventh Framework Programme (FP7/2007-2013) / ERC grant agreement n° 616905; from the European Union’s Horizon 2020 research and innovation programme under the grant agreement No 681094; from the Deutsche Forschungsge-meinschaft (DFG, German Research Foundation) – project no. 347592254 (WE 5046/4-2); from the Federal Ministry of Education and Research (BMBF) under support code 01ED2210. AP is funded by the EU-project H2020-860563 euSNN. Renzo Huber was supported by the NIMH Intramural Research Program (#ZIAMH002783) and by NWO VENI project 016.Veni.198.032 during this project.

## Data and Software availability statement

Analysis code is available on GitHub: *<*https://github.com/sdres/neurovascularCouplingVASO*>*. Raw data is available on OpenNeuro: *<*https://openneuro.org/datasets/ds004943/versions/1.0.2*>*.

## Author Contributions

According to the CRediT (Contributor Roles Taxonomy) system.

|  |  |
| --- | --- |
| Conceptualization | S.D. R.H. O.F.G. D.I. A.P. R.T. N.W. R.G. |
| Data curation | S.D. R.H. O.F.G. A.P. R.T. D.I. |
| Formal analysis | S.D. |
| Funding acquisition | R.G. N.W. |
| Investigation | S.D. R.H. R.T. R.G. A.P. O.F.G. |
| Methodology | S.D. R.H. O.F.G. D.I. R.G. |
| Project administration | S.D. R.H. R.G. |
| Resources | S.D. R.H. R.T. R.G. N.W. |
| Software | S.D. R.H. O.F.G. R.G. |
| Supervision | R.H. R.T. O.F.G. N.W. D.I. R.G. |
| Validation | S.D. |
| Visualization | S.D. R.H. O.F.G. A.P. D.I. R.G. |
| Writing - original draft | S.D. |
| Writing - review and editing | S.D. R.H. R.T. O.F.G. A.P. D.I. N.W. R.G. |

## Footnotes

1 From Gulban et al., 2022 Supplementary Figure 9: ‘The vessel diameters reported here are known to be imprecise measurements. As Duvernoy states, (I) “The diameter of cortical arteries is difficult to evaluate; the results of measurements are debatable and are a function of the pressure of injection and modifications caused by the fixation. Even in vivo … there are large variations in the diameter of cortical vessels according to physiological conditions.”; and (II) “Fixation and embedding often greatly deform veins, due to the thinness of their walls; thus, diameter measurements taken after India ink injection are of little value and are often lower than those obtained with vascular casts.”.’

## Supplementary Materials

**Figure S1:**
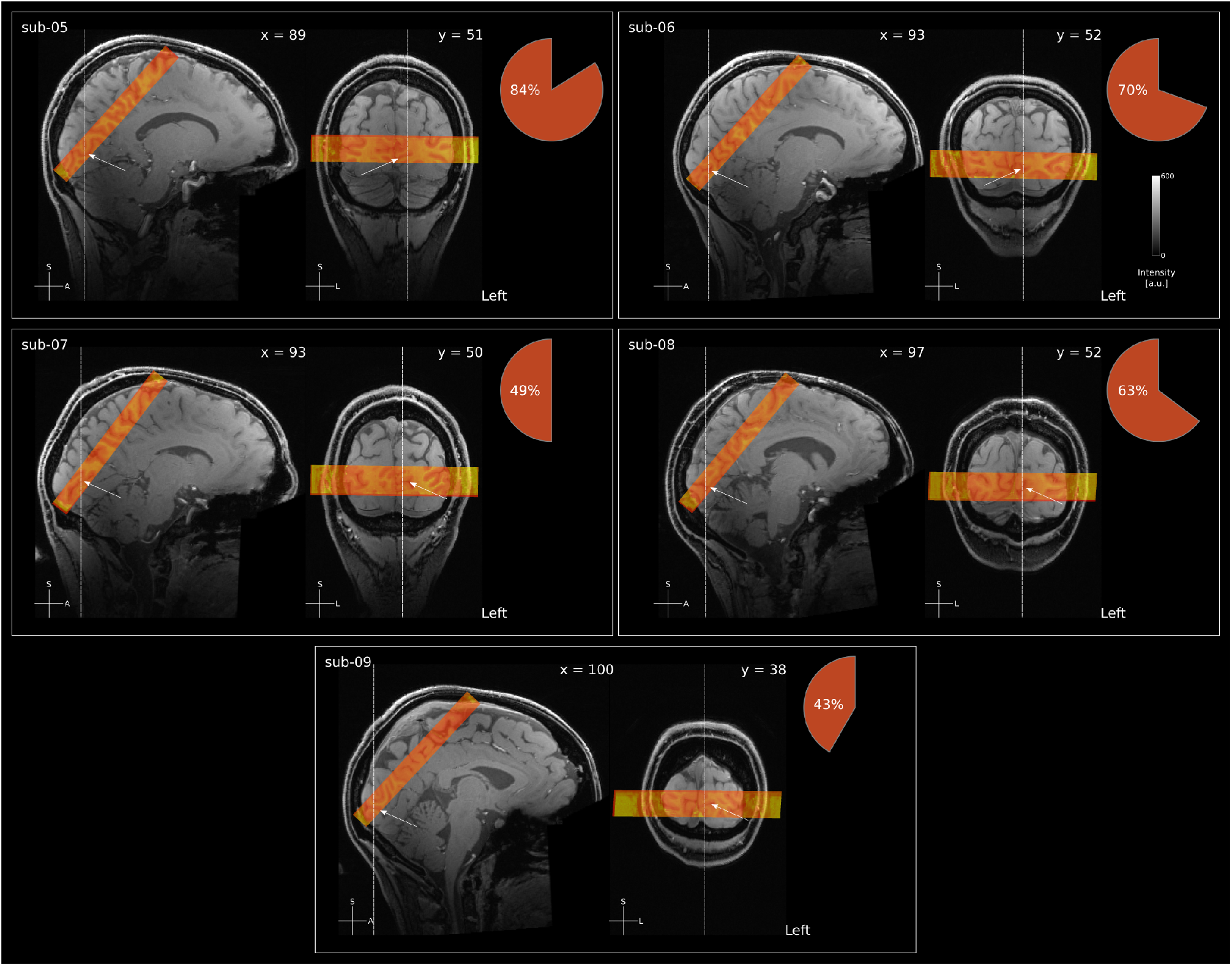
Functional data coverage for all participants. White arrows indicate location of the calcarine sulcus, based on anatomical landmarks. Because of the small FOV and number of slices, it was not always possible to include the entire calcarine sulcus without foldover. For example in participant sub-09, the sulcus folded down from the opening. The pie chart indicates an estimate of the percentage of V1 that was covered for a given participant.

**Figure S2:**
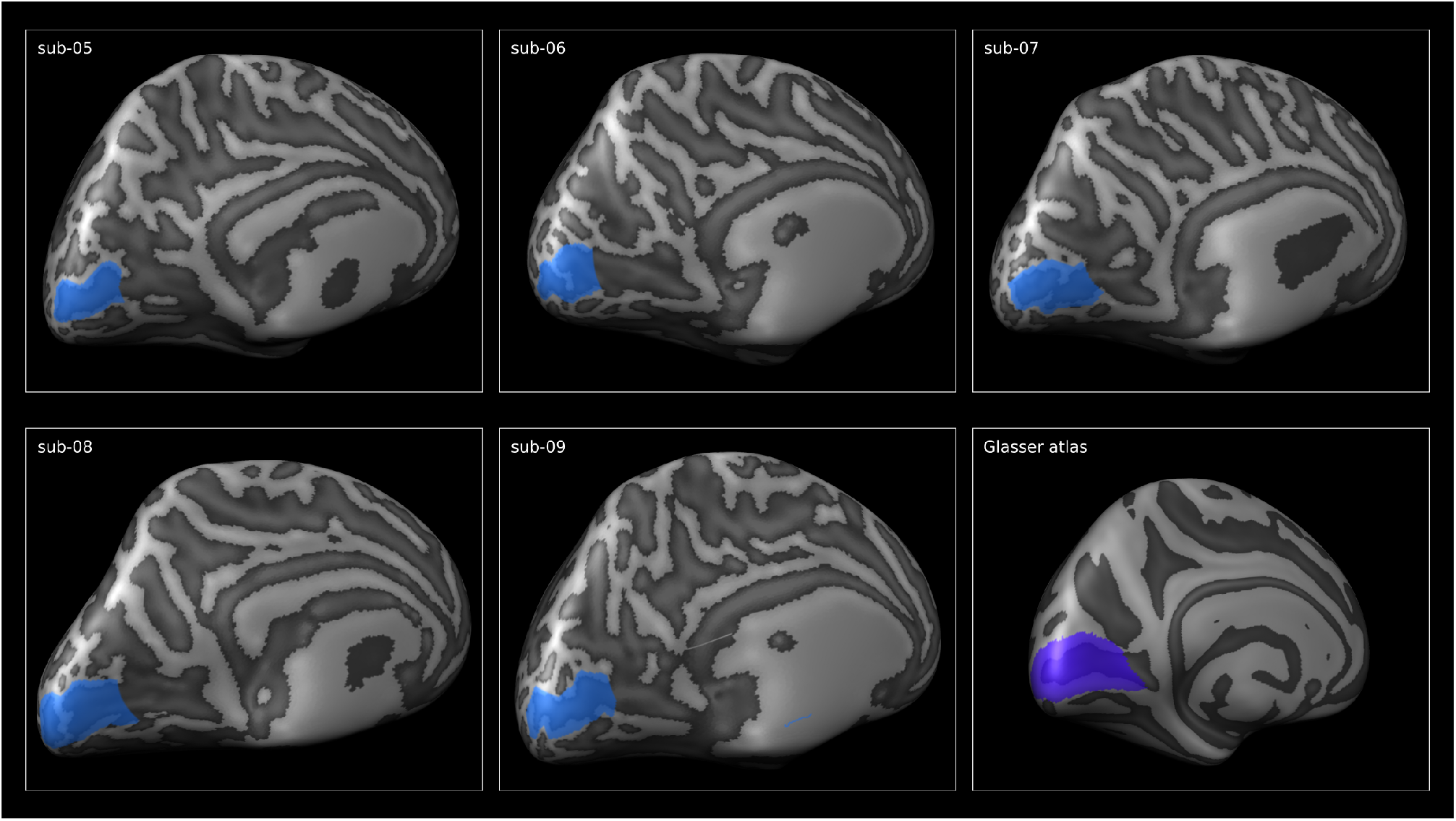
Outline of the posterior calcarine sulcus for all participants. Based on the Glasser atlas, we drew manual outlines of the calcarine sulcus (indicating V1) on the inflated cortical surface for all participants. These patches were used to estimate the overlap between the functional coverage and V1 (see **Supplementary Figure S1**). Furthermore, we used them to visually ensure that our ROIs were located in V1.

**Table S1:**
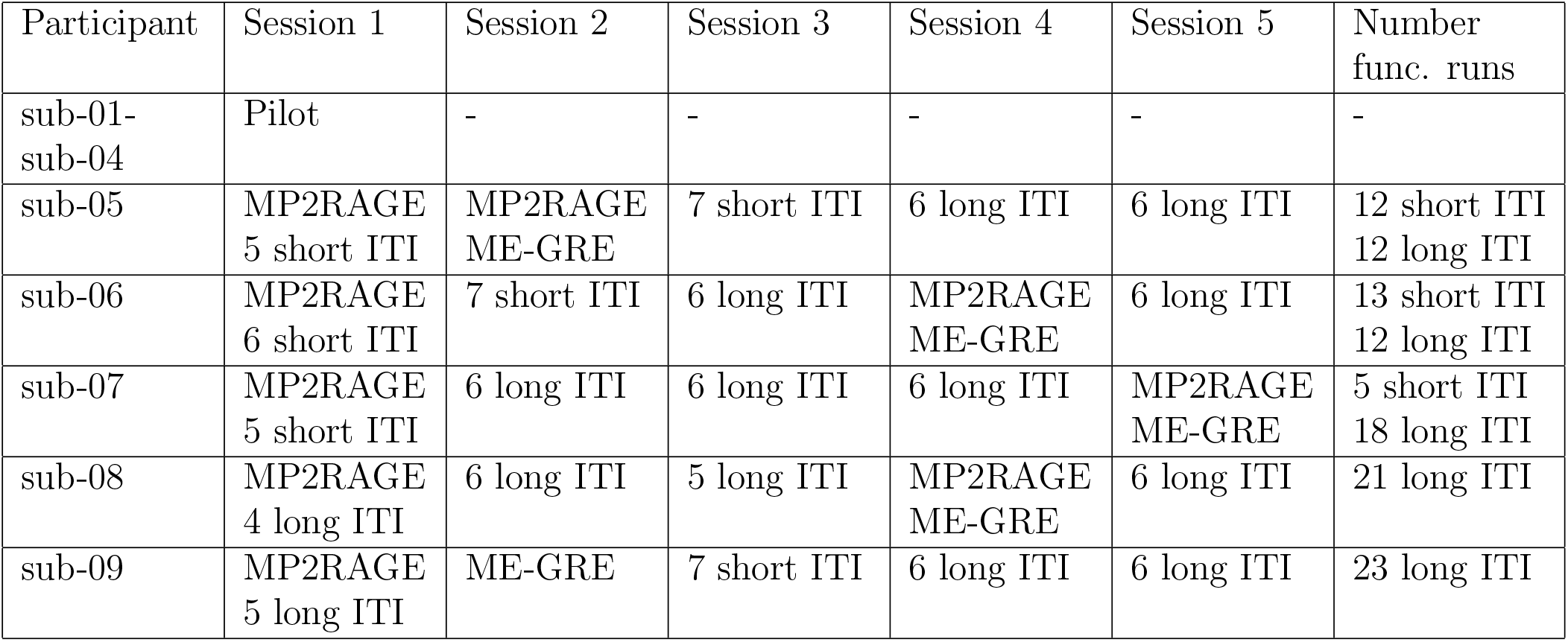
Session overview of participants.

| Participant | Session 1 | Session 2 | Session 3 | Session 4 | Session 5 | Number<br>func. runs |
| --- | --- | --- | --- | --- | --- | --- |
| sub-01-sub-04 | Pilot | - | - | - | - | - |
| sub-05 | MP2RAGE<br>5 short ITI | MP2RAGE<br>ME-GRE | 7 short ITI | 6 long ITI | 6 long ITI | 12 short ITI<br>12 long ITI |
| sub-06 | MP2RAGE<br>6 short ITI | 7 short ITI | 6 long ITI | MP2RAGE<br>ME-GRE | 6 long ITI | 13 short ITI<br>12 long ITI |
| sub-07 | MP2RAGE<br>5 short ITI | 6 long ITI | 6 long ITI | 6 long ITI | MP2RAGE<br>ME-GRE | 5 short ITI<br>18 long ITI |
| sub-08 | MP2RAGE<br>4 long ITI | 6 long ITI | 5 long ITI | MP2RAGE<br>ME-GRE | 6 long ITI | 21 long ITI |
| sub-09 | MP2RAGE<br>5 long ITI | ME-GRE | 7 short ITI | 6 long ITI | 6 long ITI | 23 long ITI |

**Figure S3:**
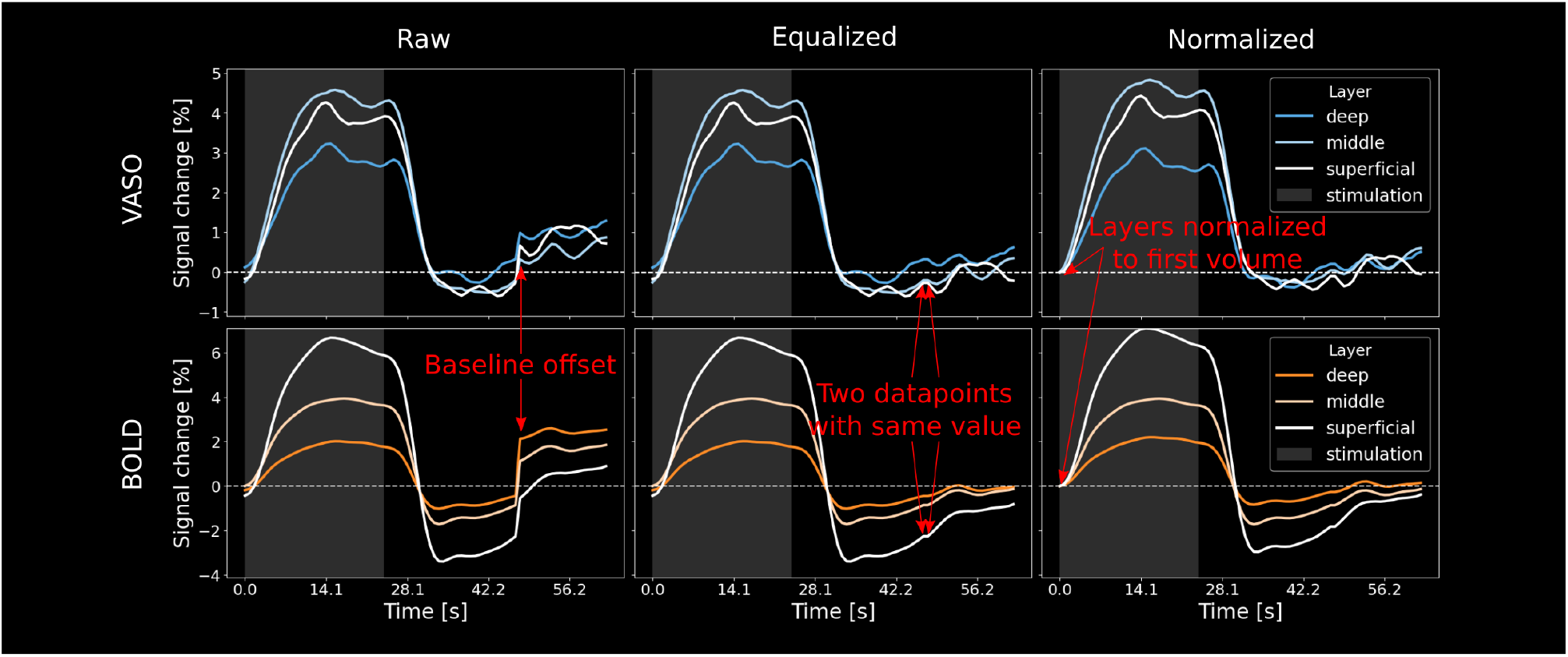
Equalization and normalization procedure. Left: VASO and BOLD ERAs for one individual participant (sub-06) in terms of raw extracted % signal change. A shift in baseline is clearly visible between sessions with short to long ITIs. Middle: Volumes with long ITIs were matched with the last time point of the short ITI sessions. This leads to two timepoints with the same value. Right: Finally, we set the first volume of each layer to 0 and adjusted all remaining volumes accordingly.

**Figure S4:**
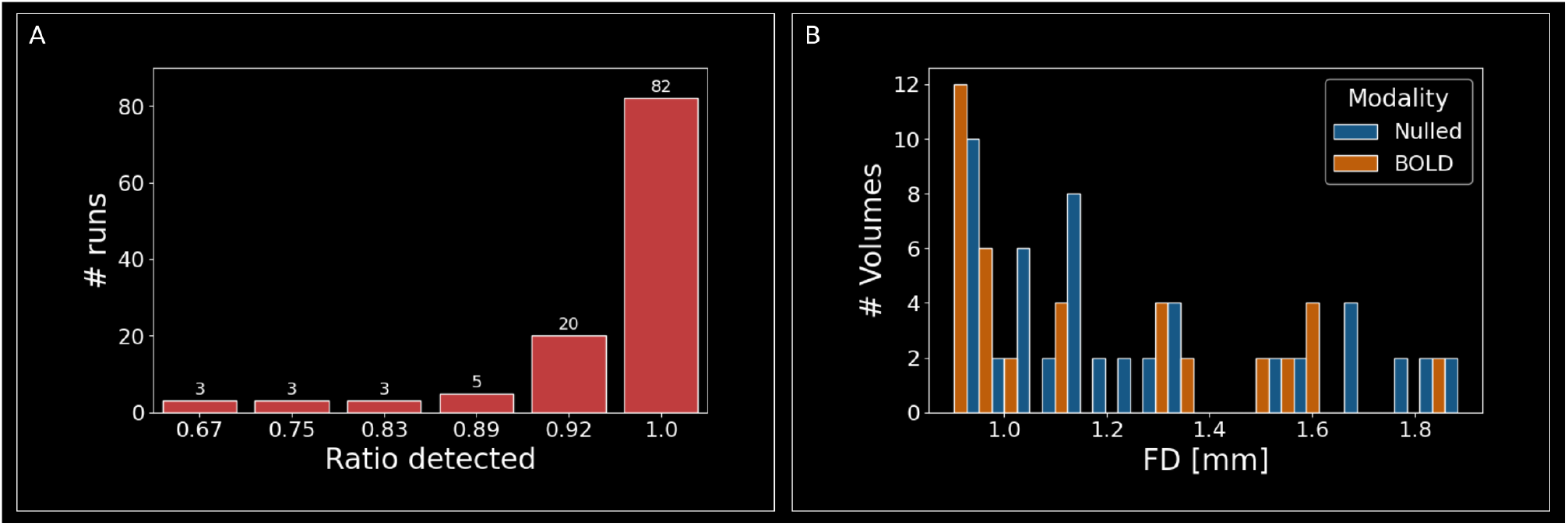
Attention-task performance and motion. **A** Number of targets detected divided by the total number of targets per run. Performance was high overall, with only a limited number of runs in which multiple targets were undetected. **B** Distribution of volumes with FD ¿ 0.9 mm across BOLD and nulled acquisitions. We acquired a total of 117088 volumes (nulled and BOLD combined). Only 92 volumes (¡0.08%) showed framewise displacements (FDs) greater than our voxel size (0.9 mm). 40 (in 16 individual runs) of those were in BOLD and 52 (in 12 individual runs) were in nulled time series and FD never exceeded 1.88 mm. We therefore did not exclude any data based on motion.

**Figure S5:**
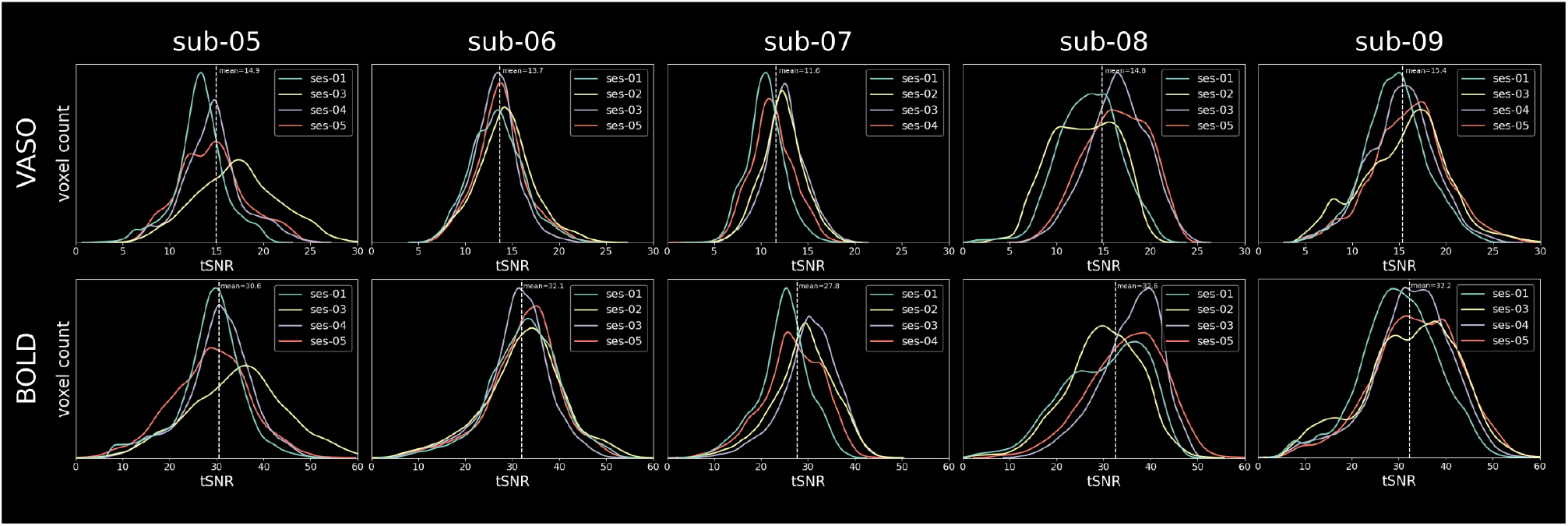
VASO and BOLD tSNR for each participant and session individually. VASO and BOLD tSNR values are mostly uniform across sessions with minor differences between participants. Values were extracted from the same regions of interest as GLM results and signal changes

**Figure S6:**
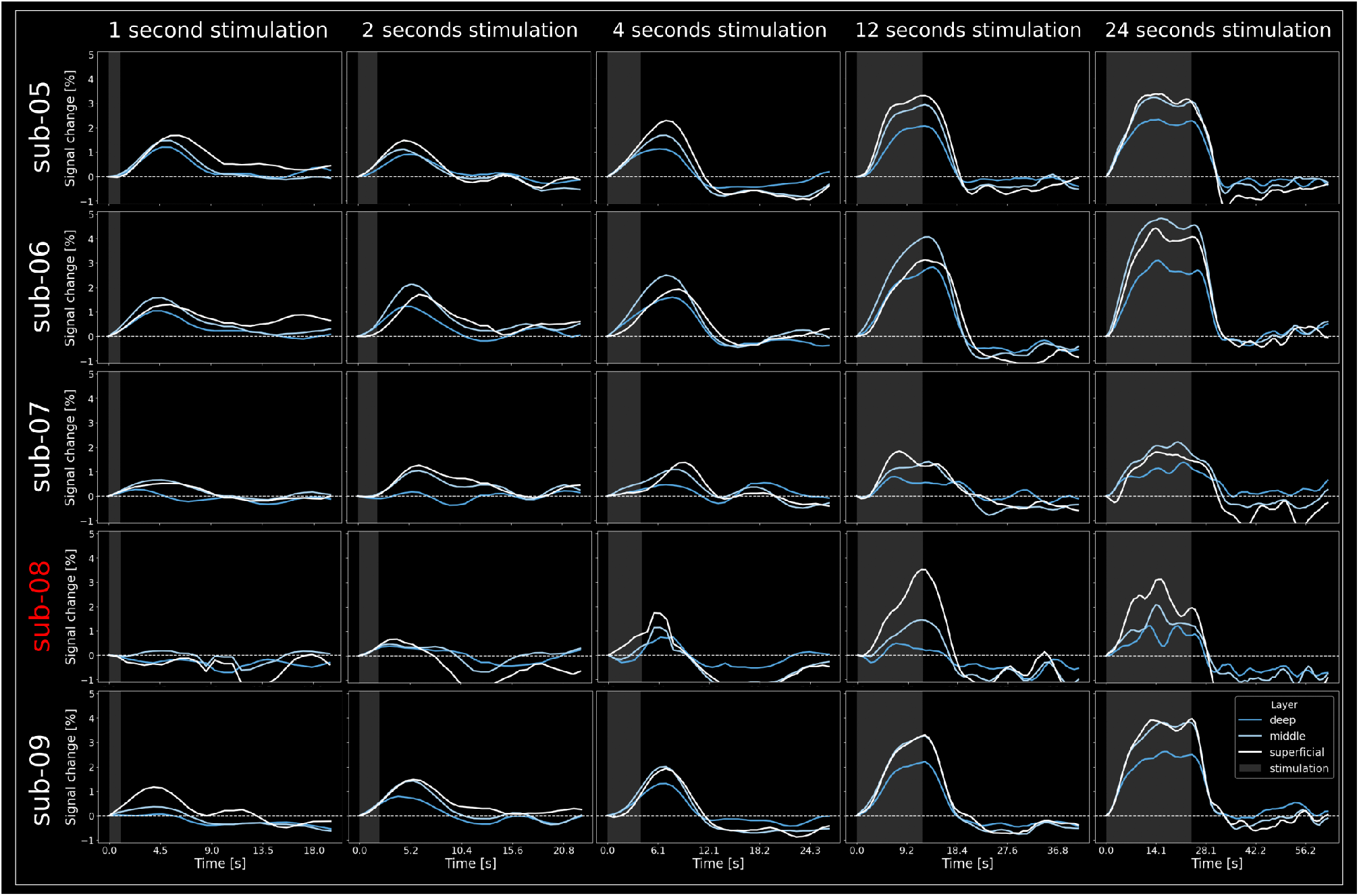
VASO results of individual participants.. Same as **Figure 3B** but for all participants individually. Data averaged across all sessions and normalized to zero to the first datapoint for each layer. Note that participant sub-08 did not show any activation in response to stimulation of 1 second and only weak activation in response to 2 second stimulation. This prohibits some of the normalization procedures conducted in this study (e.g. division by zero for data in **Figure 7**. Therefore, we excluded this participant in the group average plots.

**Figure S7:**
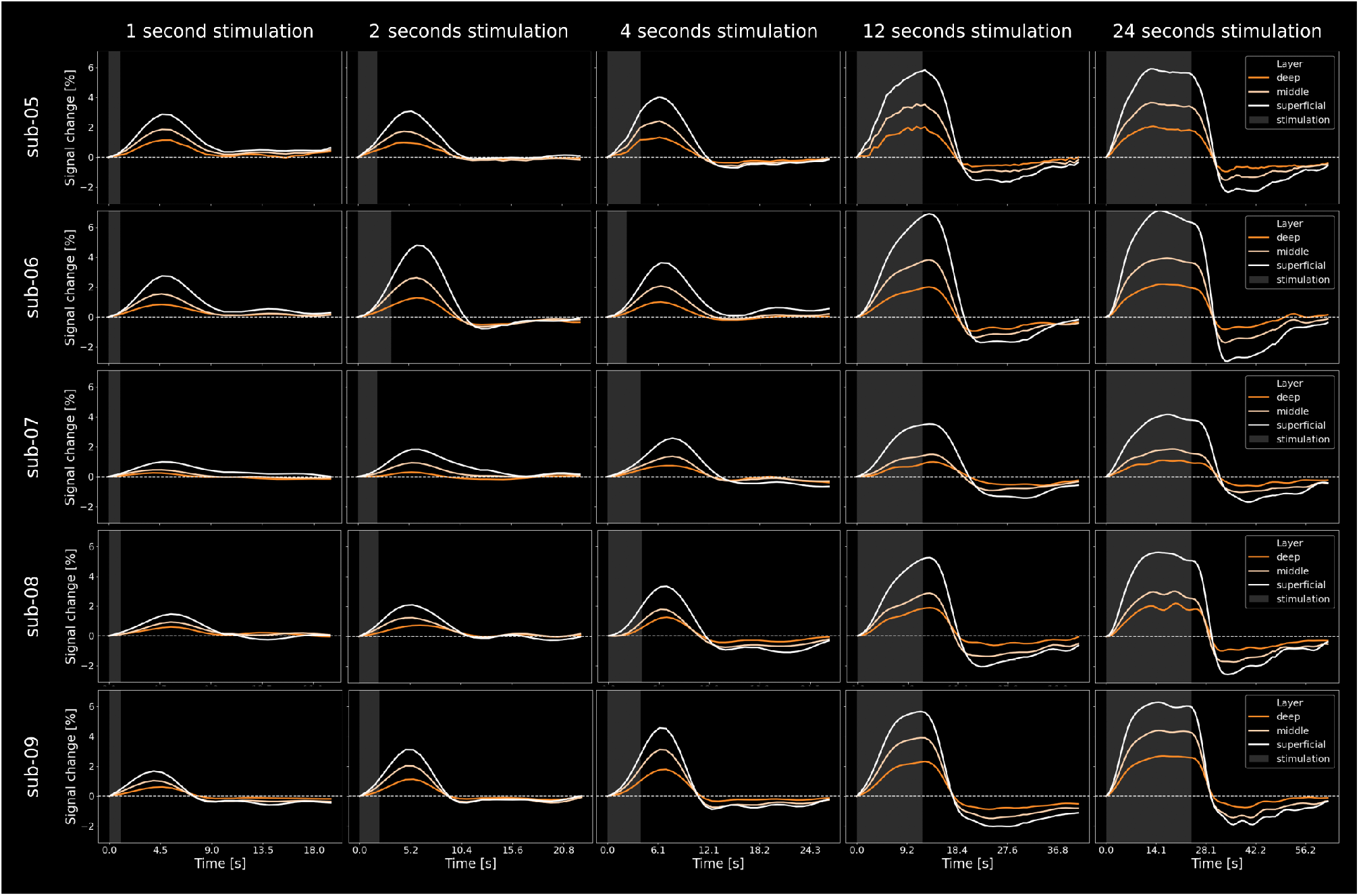
BOLD results of individual participants. BOLD results of individual participants. Same as **Figure 3B** but for all participants individually. Data averaged across all sessions and normalized to zero to the first datapoint for each layer.

**Figure S8:**
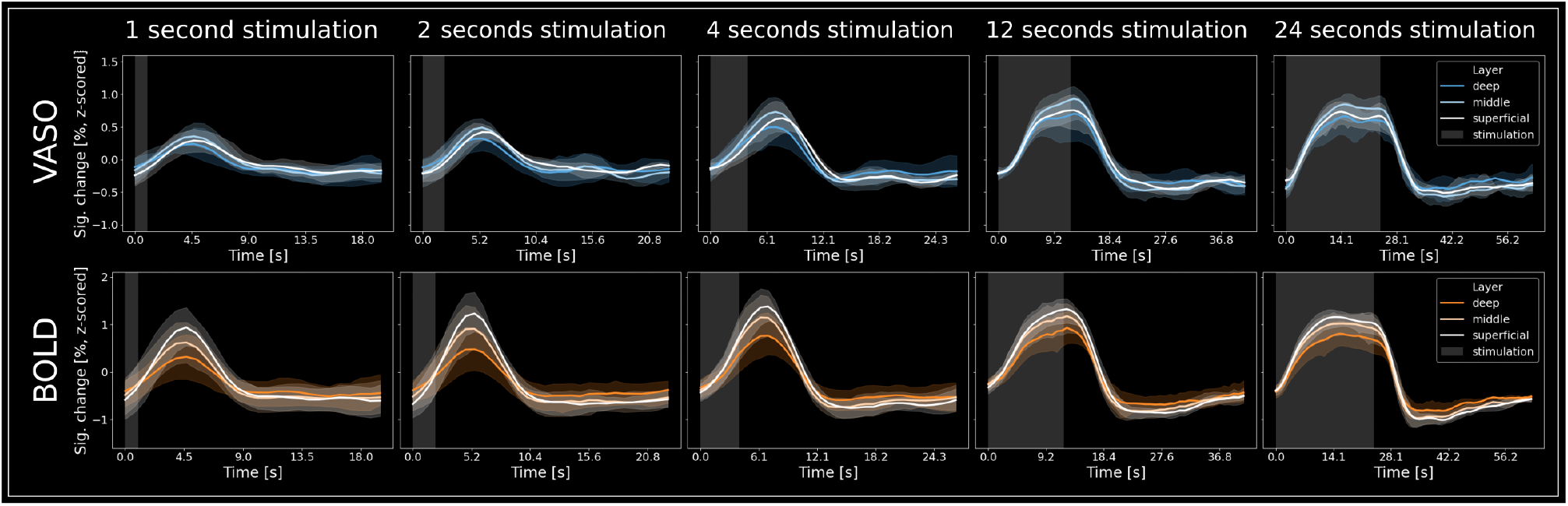
Group-level VASO and BOLD (z-scored). Same as **Figure 3B** but z-scored.

**Figure S9:**
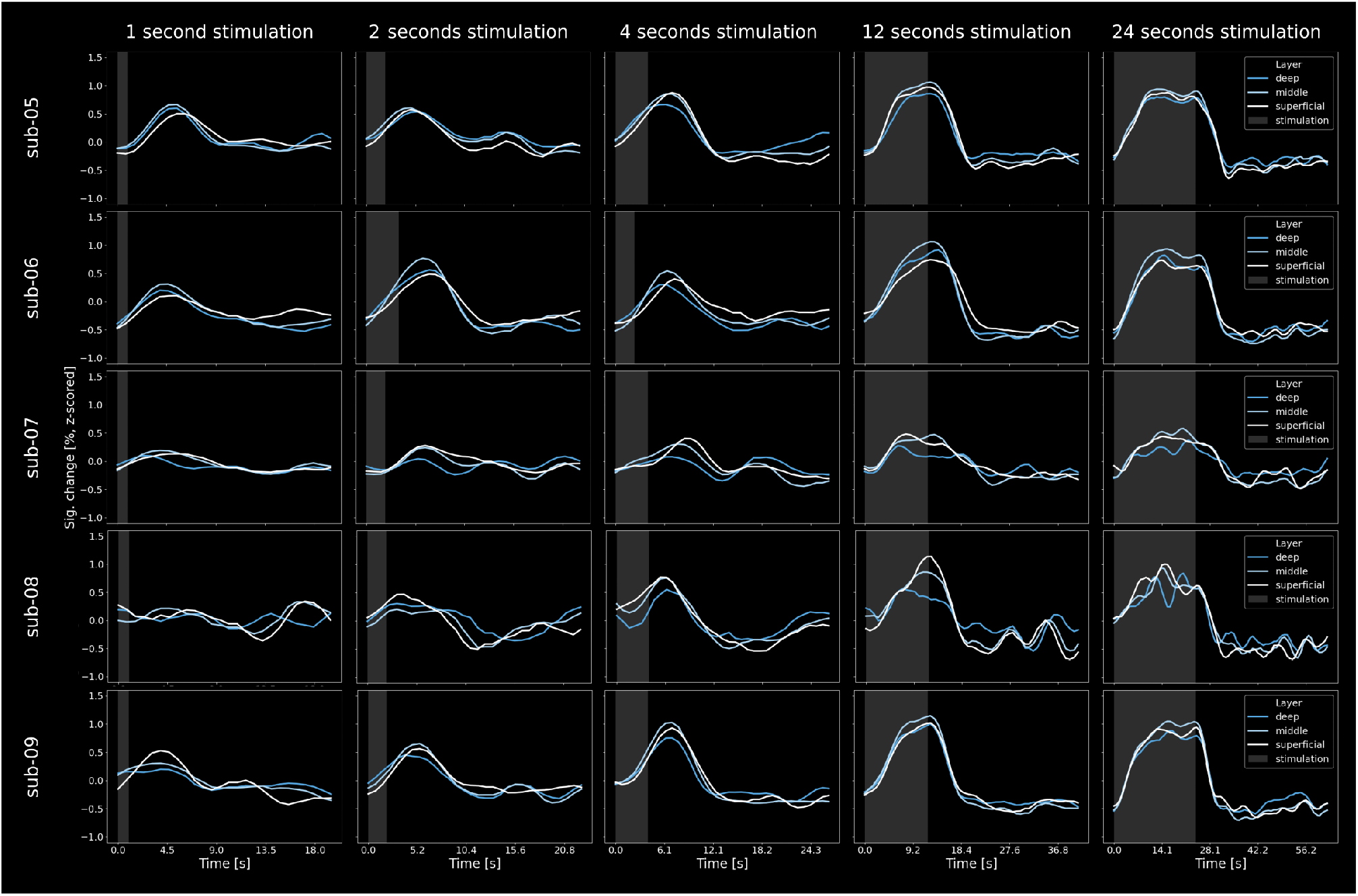
Z-scored VASO results of individual participants. Same as Figure S6 but with z-scored signal changes.

**Figure S10:**
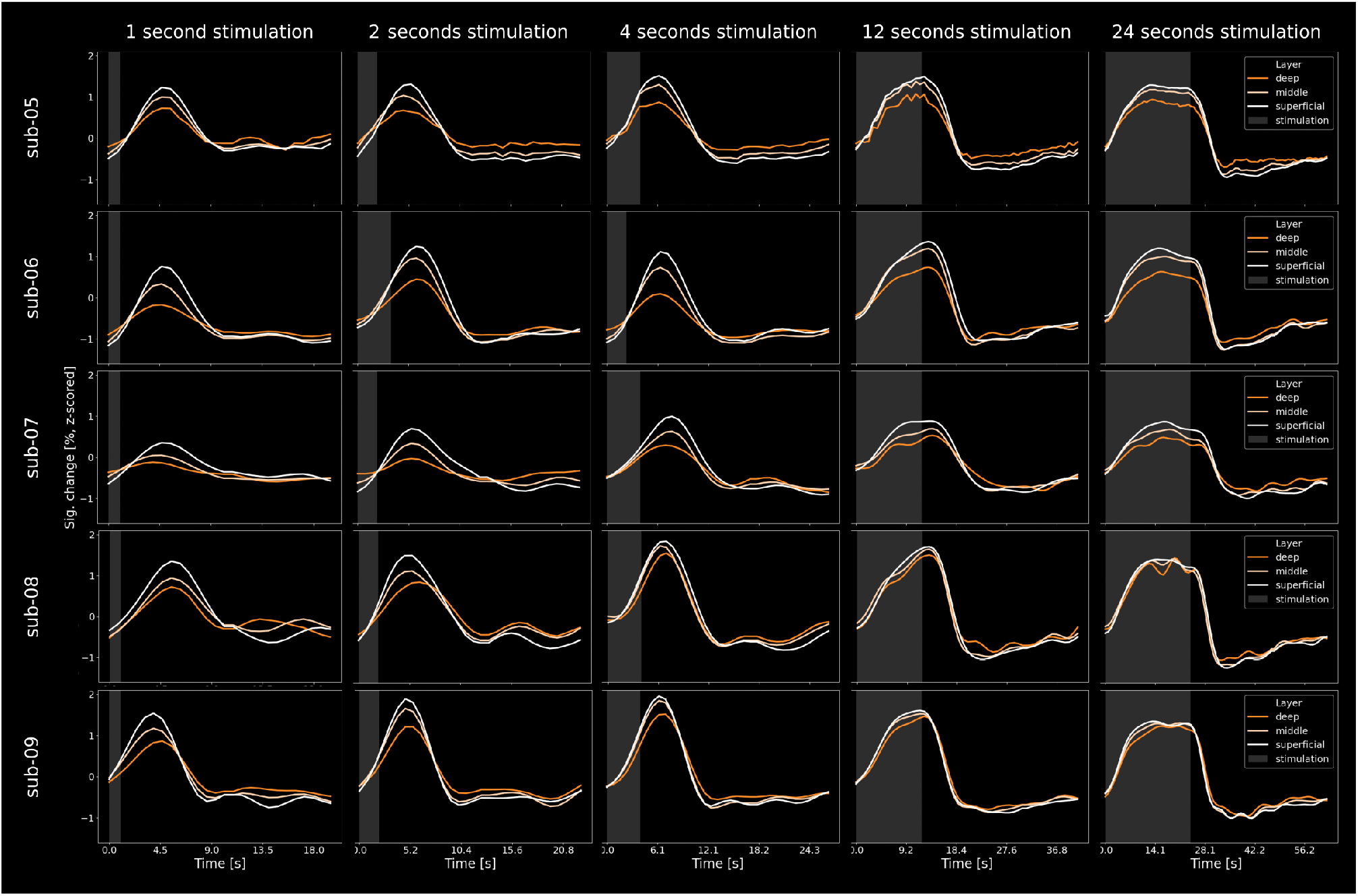
Z-scored BOLD results of individual participants. Same as Figure S7 but with z-scored signal changes.

**Figure S11:**
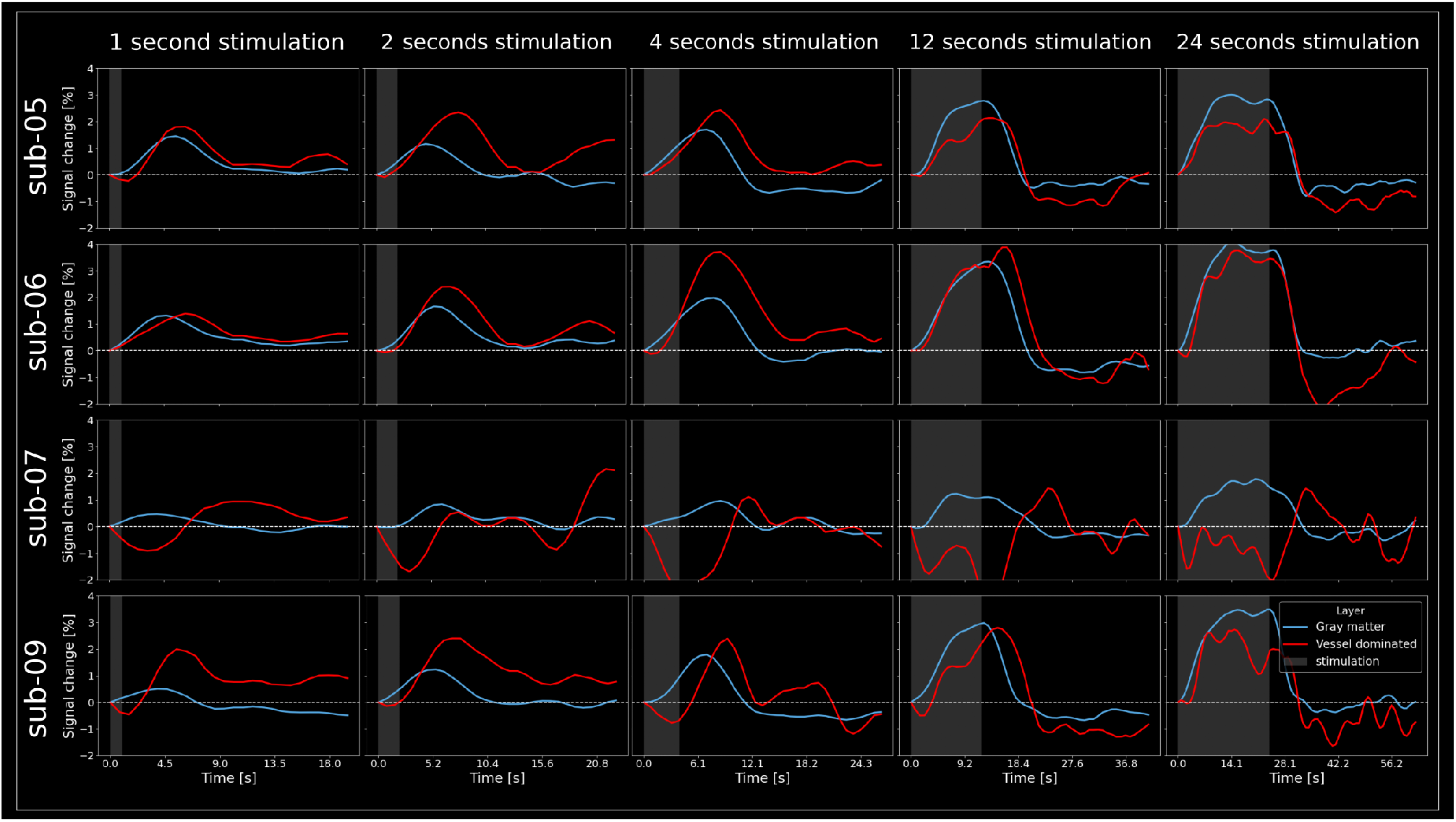
VASO responses in vessel dominated and gray matter voxels of individual participants. Supplement to **Figure 6**

**Figure S12:**
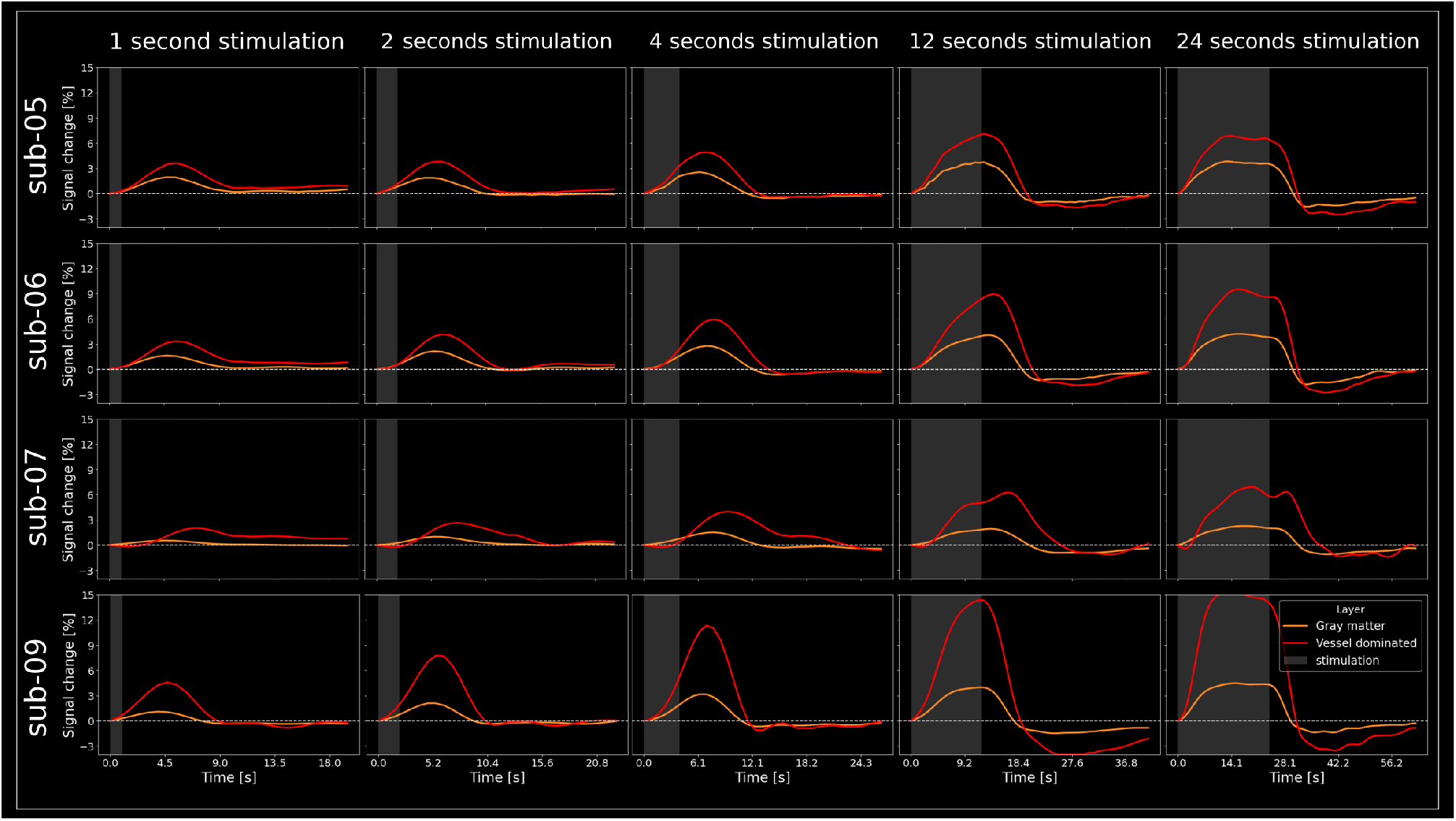
VASO responses in vessel dominated and gray matter voxels of individual participants. Supplement to **Figure 6**

## BrainVoyager pipeline for surface reconstruction

We applied the following BrainVoyager pipeline to generate inflated cortical surfaces for all participants. A video description can be found here: *<*https://www.youtube.com/watch?v=5Ik71Y_cLS8&ab_channel=ofgulban*>*

1. MP2RAGE denoising
2. Brain extraction (mask size: −10)
3. Mask smoothing (FWHM 10 mm gaussian)
4. Erode mask (2 steps)
5. Intensity inhomogeneity correction
6. Iso voxeling to 0.5 mm isotropic resolution
7. ACPC transformation
8. Set Talairach box without interpolation
9. Label ventricles as WM (using talairach definition)
10. Remove cerebellum (using talairach definition)
11. Tissue contrast enhancement using sigma filter
12. Calculate gradients file
13. Adaptive wm-gm segmentation
14. Polish
15. Estimate GM (not crucial)
16. Disconnect hemispheres
17. Run bridge removal tool (for details, see Kriegeskorte & Goebel, 2001)
18. Reconstruct surface mesh
19. Mesh smoothing
20. Mesh simplification (80k vertices)

